# Evolution with complex selection and transmission

**DOI:** 10.1101/696617

**Authors:** Sean H. Rice

**Affiliations:** Dept. of Biological Sciences, Texas Tech University

## Abstract

Inheritance is the key factor making biological evolution possible. Despite this central role, transmission is often bundled into the simplifying assumptions of evolutionary models, making it difficult to see how changes in the patterns of transmission influence evolutionary dynamics. We present a mathematical formalism for studying phenotypic evolution, under any selection regime and with any transmission rules, that clearly delineates the roles played by transmission, selection, and interactions between the two.

To illustrate the approach, we derive models in which heritability and and fitness are influenced by the same environmental factors – producing a covariation between selection and transmission. By itself, variation in heritability does not influence directional evolution. However, we show that any covariation between heritability and selection can have a substantial effect on trait evolution. Moderate differences in heritability between environments can lead to organisms adapting much more to environments with higher heritability, and can pull a population off of an “adaptive peak”. When habitat preference is allowed to evolve as well, variation in heritability between environments can lead to organisms exclusively using the environment in which heritability is highest. This effect is most pronounced when initial habitat selection is weak.

## Introduction

Evolution is influenced both by patterns of survival and reproduction, which contribute to both selection and drift, and by patterns of transmission, broadly defined to encompass the phenotypic relationship between offspring and parents (or descendants and their ancestors). With the exception of migration, all evolutionary mechanisms involve some combination of these categories. This is manifest in the fact that the most compact general equation for evolutionary change, the stochastic Price equation, represents change in mean phenotype in terms of the relationship between offspring phenotype (determined by transmission) and parental fitness.

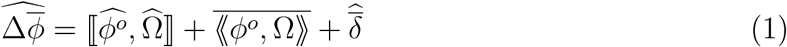

In Equation 1, *ϕ°* represents the mean phenotype of an individual’s offspring, and Ω represents relative fitness (see Table 1 for notation). ⟦, ⟧ is the frequency covariance of two values among individuals in a population, and ⟪, ⟫ is the probability covariance between two random variables in an individual (see Methods and Rice and Papadopoulos (2009) for discussion of the relationship between frequency and probability operations).

**Table 1:** Symbols and Notation

|  |  |
| --- | --- |
| $\bar{X}$ | Average value of $X$ across its frequency distribution |
| $\widehat{A}$ or $E(A)$ | Expected value of random variable $A$ |
| $\llbracket^2 X \rrbracket$ | Variance in the value of $X$, across its frequency distribution |
| $\llbracket^j X \rrbracket$ | $j^{th}$ central moment of the frequency distribution of $X$ |
| $\langle^2 A \rangle$ | Variance in random variable $A$ across its probability distribution |
| $\langle^j A \rangle$ | $j^{th}$ central moment of the probability distribution of random variable $A$ |
| $\llbracket X, Y \rrbracket$ | Frequency covariance of $X$ and $Y$ |
| $\langle A, B \rangle$ | Probability covariance of random variables $A$ and $B$ |
| $\llbracket_b^a \rrbracket$ | Simple regression of $a$ on $b$. |
| $w$ | Absolute fitness (number of descendants after one generation) |
| $\Omega$ | Relative fitness ($\Omega = \frac{w}{\bar{w}}$), conditional on $\bar{w} \neq 0$ |
| $\phi$ | Phenotype of an individual |
| $\phi^o$ | Average phenotype of an individual's offspring |
| $\tilde{x}(\phi)$ | Average value of $x$ among individuals that share the same phenotypic value |

If we specify the patterns of reproduction and transmission for a particular kind of system, then Equation 1 gives us a model of evolution in that system. For example, interpreting *ϕ* as allelic type and assuming that transmission is Mendelian yields equations for change in allele frequency from classical one locus population genetics. Similarly, treating *ϕ* as a continuous trait, assuming that *ϕ°* is a linear function of *ϕ*, and assuming that population size is constant and large yields the “breeder’s equation” from quantitative genetics (Rice, 2004).

Though various other transmission rules have been studied – such as two loci with recombination, maternal inheritance, etc. – there remain systems that are not well approximated by any of these special cases. For example, Figure 1 shows two examples of nonlinear relationships between offspring and parents for continuous traits.

**Figure 1:**
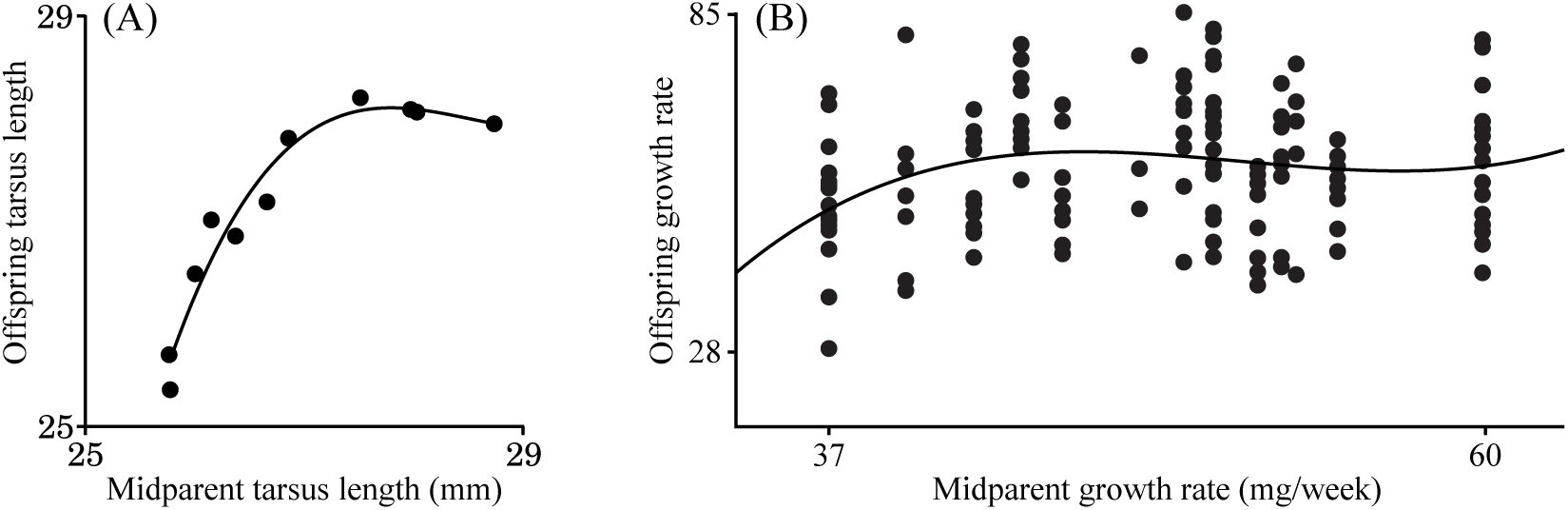
Examples of nonlinear transmission: (A) Relation between offspring tarsus length and the midparent value in hybrid buntings. Data from Ryan (2001). (B) Relation between offspring and midparent growth rate in earthworms. Data from McElroy and Diehl (2001)

Beyond the examples in Figure 1, a number of other studies have shown that the relationship between offspring and parents can be significantly nonlinear (Beardsley et al., 1950; Nishida, 1972; Meyer and Enfield, 1975; Shimizu and Awata, 1979; Gimelfarb, 1986; Gifford and Barker, 1991; Gimelfarb and Willis, 1994; Fuerst-Waltl et al., 1998), and that this non-linearity can influence the response to selection (Gimelfarb and Willis, 1994). In cases like these, we can not get away with using a single parameter, such as additive genetic variance (the covariance between offspring and midparent phenotypes) or heritability (the regression of offspring on parents), to describe transmission.

As with the case of transmission, fitness functions have generally been modeled on a few special cases (linear, Gaussian, etc.). Once again, though these special cases have yielded valuable models, they are likely to be poor approximations to many natural systems. We thus seek a way to combine selection and transmission that does not limit the kinds of fitness functions and transmission rules that we can study.

The principal mechanistic link between offspring phenotype (*ϕ°*) and parental fitness (Ω) is the fact that they are both influenced by parental phenotype (*ϕ*). We thus seek a way to write both offspring phenotype and parental fitness in terms of the phenotypes of parents. One way to do this would be to write both *ϕ°* and Ω in terms of a nonlinear function of phenotype. Unfortunately, in standard polynomial regression, the regression coefficients are functions of the degree of the polynomial that we choose to fit, which is arbitrary. As an example, the first, second, and third order approximations to the earthworm data in Figure 1(B) are as follows:

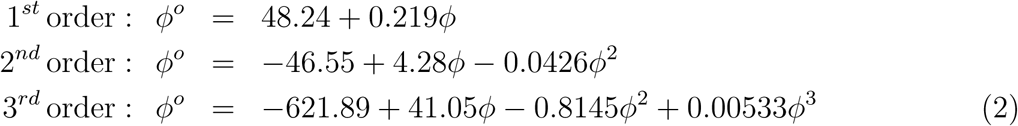

The coefficient of *ϕ* in the linear function (0.219) is the “heritability” of this trait (defined here as the slope of the linear regression of offspring on midparent phenotype). As we fit higher order polynomials to the data, however, the coefficient of the linear term changes by roughly an order of magnitude for each new term that we add to the function. This is because different powers of *ϕ* tend to be correlated with one another.

The fact that all of the regression coefficients change as we consider higher order polynomials poses two significant problems. First, the heritability does not appear when we apply even a second order regression – making it difficult to directly compare the more complex model with the linear case (the same problem arises when we use the additive genetic variance instead of heritability). Second, and more troubling, because the regression coefficients change as we change the degree of the polynomial – and there is no clear “natural” degree of polynomial that we should favor – the coefficients in a standard polynomial regression have no distinct biological meaning.

What we need to do is represent variables like offspring phenotype or fitness in terms of functions of *ϕ* that capture nonlinearity, but are not themselves correlated with one another. Ideally, these functions would include *ϕ* itself, so that traditional concepts of heritability and selection differentials would emerge naturally, along with other terms that capture more complex relationships. Below we show how this can be done – for offspring phenotype, fitness, or any other value – using a set of orthogonal polynomials defined by the population itself (see the Supplementary Material for a general description of orthogonal polynomials).

Orthogonal polynomials are constructed so that each is independent of (orthogonal to) all of the others. For functions, orthogonality is defined relative to a “weight function”. There are a number of sets of classical orthogonal polynomials, each defined with respect to a different weight function. For example, the Legendre polynomials used to study function valued traits such as continuous growth processes (Kirkpatrick et al., 1990; Meyer and Kirkpatrick, 2005; Pletcher and Geyer, 1999; Stinchcombe et al., 2012) have as a weight function a uniform distribution on a fixed interval, whereas the Hermite polynomials have a normal distribution as their weight function. The key to our approach is to use the population itself as the weight function.

If there are *n* distinct phenotypic values present in the population, then projecting a variable onto the polynomials up to ℙ^[*n-*1]^ yields a curve that passes through the mean value of the variable for each phenotypic value (Figure 2). If each individual has a unique value of *ϕ*, then this curve passes exactly through each point.

**Figure 2:**
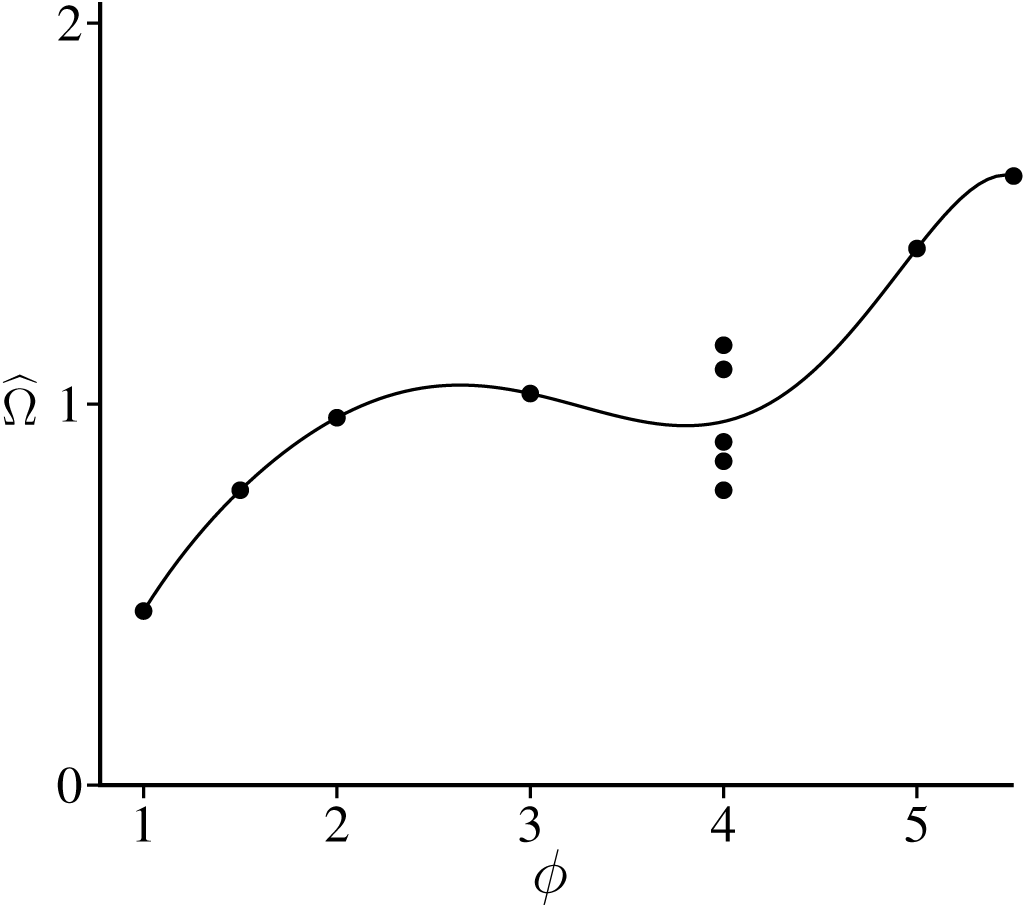
Sum of projections of 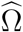 onto orthogonal polynomials: In this case there are 7 distinct values of ϕ, one of which is shared by multiple individuals. The sum of projections of 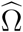 onto ℙ^[0]^ through ℙ^[6]^ yields a curve that passes exactly through points corresponding to individuals with unique phenotypic values, and through the mean when multiple individuals share the same value of ϕ.

### Offspring phenotype

We will use 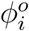 to denote the average phenotype among offspring (or descendants) of individual *i* (an “individual” can be any entity that leaves descendants; if that entity is a mated pair, then we often use the “midparent” value as the parental phenotype). To represent *ϕ°* in the space of orthogonal polynomials, *𝒫*, we project it separately onto each of the different ℙ functions. The projection on ℙ ^[0]^ is just the mean, 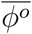, and for higher order functions (⩾1) the projection is just the simple regression of *ϕ°* on that function. Defining 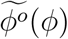 as the mean value of *ϕ °* among individuals with phenotype *ϕ*, we get:

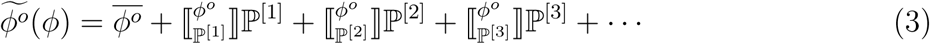

Prior to reproduction, offspring phenotype is a random variable – meaning that for each individual (or mated pair) there is a distribution of possible values of *ϕ°*. Given that the P^[*n*]^ terms are not random variables (since they are functions of the phenotypes currently present in the population), the expected value of 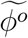 is given by:

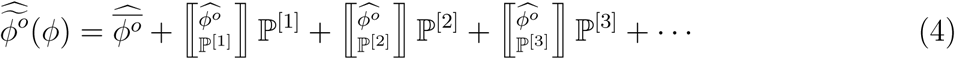

(see Methods).

To illustrate this approach, consider the data for earthworm growth in Figure 1(B). Substituting the moments of the distribution of parent phenotypes from this dataset into Equations 25 yields the following polynomials:

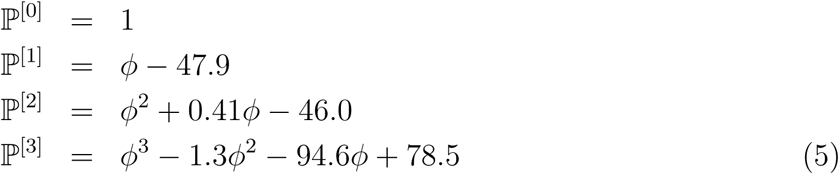

Projecting the offspring phenotype data in Figure 1(B) into the space defined by the polynomials in Equations 5 yields:

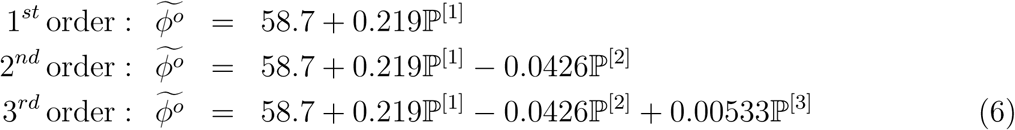

Note that the coefficients in Equations 6 do not change as we add higher order terms. Therefore, unlike the coefficients in Equations 2, we can assign them consistent biological interpretations. The first order coefficient, 0.219, captures the linear relationship between offspring and parents (*i.e.* the heritability; multiplying this by ⟦^2^*ϕ*⟧ yields the additive genetic variance), and appears no matter how many terms we add. The second order coefficient, −0.0426, measures the quadratic curvature of the offspring-parent relationship, and higher order terms capture increasingly complex components of transmission. We can still retrieve the best fit function from the projection of the points into the space *𝒫* by substituting the polynomials in Equations 5 into the ℙ^[*i*]^ terms in Equations 6.

This approach to quantifying the relationship between offspring and parents requires no simplifying assumptions about the genetic basis of the trait, nor about the distribution of residuals. It gives us a very general way to write offspring phenotype (*ϕ°*) in terms of (mid)parent phenotype (*ϕ*). In order to derive a useful general theory of trait evolution from Equation 1, we need to do the same thing with fitness.

### Fitness

We can project relative fitness (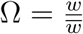, conditional on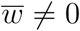) into the population space *𝒫* just as we did with offspring phenotype, yielding a function that passes through the mean value of Ω for each value of *ϕ* (denoted 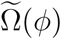). Since Ω is the ratio of individual fitness to mean population fitness, it is always the case that 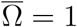, so we have:

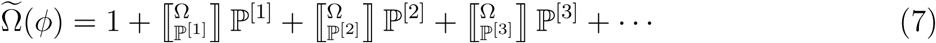

Figure 3 shows how the regression coefficients of relative fitness on the various orthogonal polynomials capture the shape of the relative fitness function. Though the function P^[2]^ is different in the examples in Figure 3A and 3B (because the phenotype distributions are different), the regressions on fitness on ℙ^[2]^ are the same, reflecting the fact that the fitness “landscape” has the same shape in both cases.

Figure 3 also illustrates the fact that the common view that the curvature of the fitness function determines the degree of stabilizing or disruptive selection (Lande and Arnold, 1983; Rice, 2004) is, in general, not true. If we consider only the effects of selection (ignoring processes, such as mutation and recombination, that are part of the process of reproduction), the expected change in phenotypic variance is then given by (see Methods):

**Figure 3:**
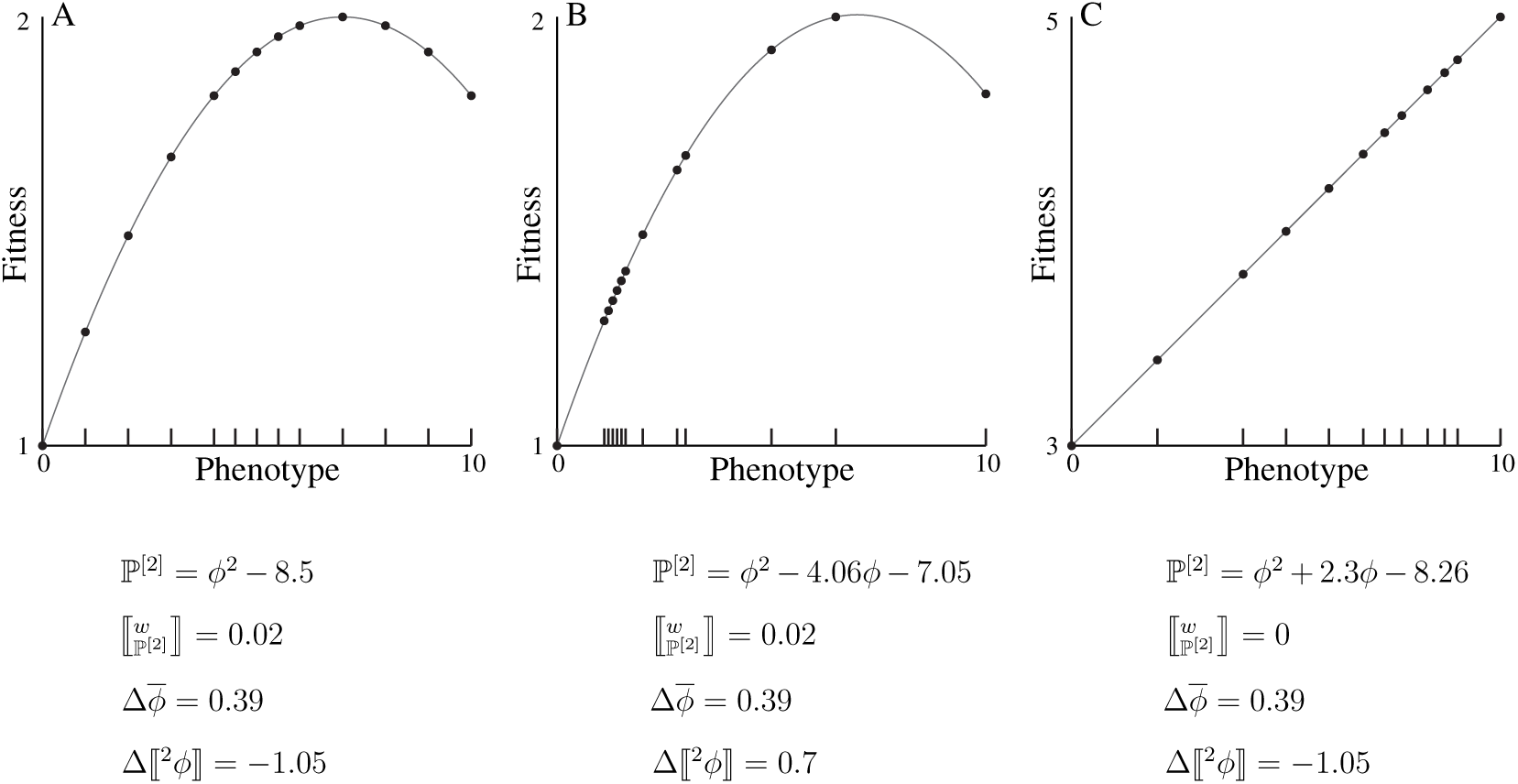
Shape of the Fitness function and Mode of Selection. Fitness as a function of phenotype for three hypothetical populations. The phenotype distributions are shown by the ticks along the horizontal axis, each tick being an individual. The dots show the fitness value for each individual.

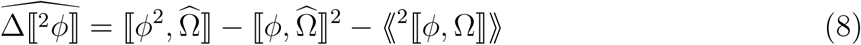

Equation 8 shows that the change in phenotypic variance is influenced by the covariance between expected relative fitness and *ϕ*^2^, which can be nonzero even when the fitness landscape is not curved, as shown by the example in Figure 3C. (Regarding the other two terms in Equation 8, 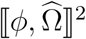 is the squared expected selection differential, and ⟪^2^⟦ *ϕ*, Ω ⟧⟫ is the variance in change due to drift.)

## Evolution of a single trait

As noted in the introduction, interpretation of Equation 1 is complicated by the fact that it combines the phenotypes of offspring (*ϕ°*) with the fitness of parents (Ω). In order to write the equation in terms of the relationship between parental phenotype and parental fitness, we substitute the series in Equation 3 for 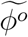 in Equation 1 yielding:

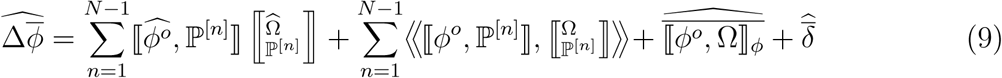

(Where *N* is population size. See Methods for derivation.)

If we consider only the first term in the first summation in Equation 9, 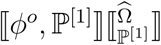we have the standard quantitative genetic equation for change in a single trait. Here, transmission is captured by covariances of expected offspring phenotype with parent phenotype, ⟦ *ϕ°*, ℙ^[1]^ ⟧, corresponding to the “additive genetic variance”, and selection is captured by regression of relative fitness on phenotype.

The 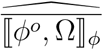 term captures change resulting from relationships between relative fitness and offspring phenotype among individuals who share the same value of *ϕ* (and thus have the same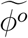), such as the 5 individuals with *ϕ* = 4 in Figure 2. If only one individual has a particular phenotypic value, then [ *ϕ°*, Ω]]_*ϕ*_ = 0 for that phenotype. If *ϕ* is a continuous variable, then each individual in a finite population is expected to have a unique value of *ϕ*, in which case we can ignore 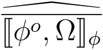 altogether.

Equation 9 is written in the form preferred by evolutionary quantitative genetics (Houle, 1992; Hansen et al., 2011), capturing transmission with covariances and selection with regressions. Alternately, we could group terms as in the “breeder’s equation”:

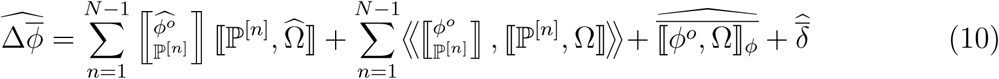

Equation 10 is essentially a generalized stochastic univariate “breeder’s equation”, since transmission is captured by regressions of offspring on parents (the first order term being heritability), and selection is captured by covariances between relative fitness (Ω) and phenotype (the first order term being the selection differential).

Note that the terms in Equations 9 and 10 always involve a covariance and a regression. If we choose to capture transmission with heritability (a regression), then we are constrained to capture selection with a selection differential (a covariance). Conversely, if transmission is captured by additive genetic variance (actually a covariance), then selection emerges as a regression of fitness on phenotype. The two forms of the equation are equivalent; they differ only in whether the phenotypic variance is included in the transmission term or the selection term.

Equations 9 and 10 do not require the relationship between offspring and parents to be fixed. In fact, the additive genetic variance and heritability are likely to change as the population evolves or, as in the examples presented below, the environment changes. If the actual relationship between offspring and parents is not linear, then both the additive genetic variance and the heritability will change as the mean phenotype changes. One advantage of the orthogonal polynomial approach is that it allows us to distinguish cases in which these values change only because of changes in the distribution of phenotypes in the population (in which case the nonlinear function relating offspring and parents would not change from one generation to the next), from cases in which the biology of transmission or development is itself evolving.

Both offspring phenotype (*ϕ°*) and relative fitness (Ω) are random variables, meaning that each individual has a distribution of possible values of *ϕ°* and Ω. This means that additive genetic variance-like terms (⟦*ϕ*^*o*^, ℙ^[*i*]^ ⟧) can covary with selection 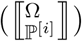. The terms in the second summation on the righthand side of Equation 9, 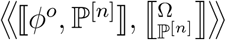, capture this probability covariance between transmission and selection. One situation in which we should expect covariation between selection and transmission is when both selection and transmission are influenced by a variable environment. We consider a simple case of this below.

### Example: Covariation between selection and transmission

A number of studies have demonstrated that a trait may have different heritabilities in different environments (Hoffmann and Merila, 1999; Merila and Sheldon, 2001; Charmantier and Garant, 2005; Ardia and Rice, 2006; Husby et al., 2011; Saastamoinen et al., 2013; Kole and Saha, 2013; Casco et al., 2014). Observed heritability values vary from slightly negative to greater than 1 (Konarzewski et al., 2005; Merila and Sheldon, 2001), with values between 0.1 and 0.5 being common. (Values outside of the range [0, 1] are possible when heritability is defined as the regression of expected offspring phenotype on midparent phenotype, as we are doing here).

Given this influence of environment on transmission, and the fact that selection regimes are likely to differ in different environments, it is not surprising that selection and heritability are often correlated (Charmantier and Garant, 2005; Wilson et al., 2006; Husby et al., 2011). In this section, we investigate some consequences of such transmission-selection covariance in a simple example with two environments.

Note that, in order to focus on a particular process, we will now introduce simplifying assumptions. This is an important point: Equations 9 and 10 are very general, being derived from a few basic axioms and thus encompassing everything that could drive change in mean phenotype. This generality comes at a price; when expanded, Equation 9 contains 2*N* terms (where *N* is population size) Our subsequent results in this section are thus not axiomatic. One point that we hope to make is that one of the values of axiomatic theories is that they can serve as formulas for producing special case models.

Consider a case in which organisms encounter one of two different environments with equal probability (we will allow the probability to evolve later), each environment imposing a different heritability. For simplicity, we consider a case in which transmission is linear in both environments, but with different slopes, and that trait values are normally distributed with fixed variance. In such a case, we need only consider the first order terms in Equations 9 or 10. If the trait is truly continuous, then no two individuals have the exact same value of *ϕ*, so 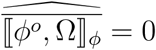, and we can write:

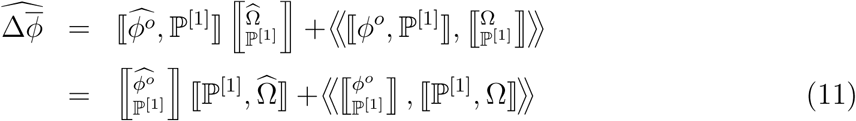

Equation 11 presents both ways of writing our result, either in terms of the “additive genetic variance” 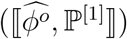 or heritability 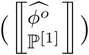. In the examples that follow, we will generally refer to heritabilities, since this is the value that is most often estimated from natural populations and in the lab. Because the variance in phenotype is the same (⟦^2^*ϕ* ⟧ = 0.001) in each example, the additive genetic variance is always 0.001 times the heritability.

In addition to imposing different heritabilities, we also stipulate that the two environments impose different selection regimes. In Environment 0, the optimal phenotypic value is *ϕ* = 0, while *ϕ* = 1 is optimal in Environment 1 (Figure 4A). Specifically, we use the following functions for fitness within the two environments:

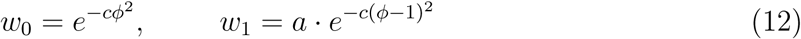

**Figure 4:**
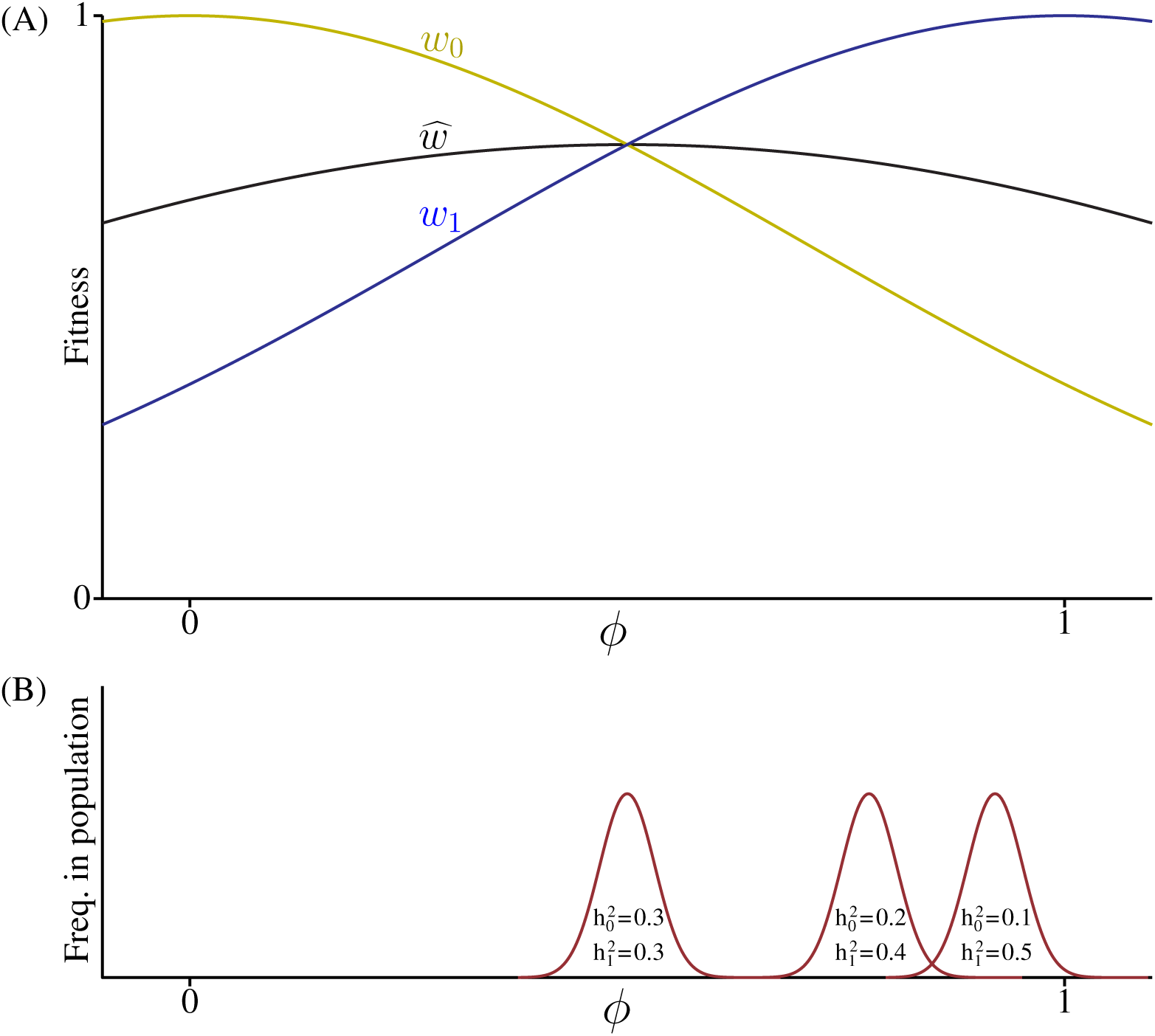
Covariation between transmission and selection. (A): Fitness in Environment 0 (*w*_0_), in Environment 1 (*w*_1_), and the expected fitness 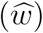 for an individual that has a probability of 0.5 of finding itself in either environment. (B): Equilibrium phenotype distributions for a population with ⟦^2^*ϕ* ⟧ = 0.001 and with different combinations of heritabilities in the two environments.

Tuning the variables *a* and *c* in Equations 12 changes the shape of the overall expected fitness function.

In Figure 4, the fitness functions are chosen so that there is a single maximum of expected fitness at 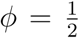(*c* = 1, *a* = 1 in Equation 12). Figure 4B shows the equilibrium phenotype distributions corresponding to different combinations of heritability values for the two environments (calculated by setting 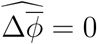 in Equation 11, see Methods).

When the trait, *ϕ*, has the same heritability in both environments, the equilibrium phenotype distribution is centered on the maximum of expected fitness 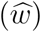. This is the outcome that would be predicted by quantitative genetic models. (To see this, note that in this case 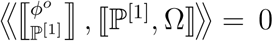, so the change in mean phenotype is given by 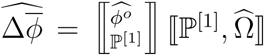 which is just the standard “breeder’s equation”.) 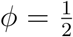 would also be the ESS state under evolutionary game theory.

When the transmission of *ϕ* is different in the different environments, however, the equilibrium changes. 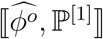 is greater in Environment 1, then for 0 *< ϕ <* 1 there is a positive probability covariance between transmission and selection 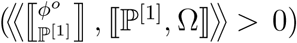 This shifts the equilibrium towards higher values of *ϕ*. Even for the case of 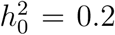 and 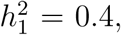, the population ends up much better adapted to Environment 1, even though individuals encounter Environment 0 half of the time and have appreciable heritability there. The case of 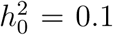 and 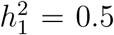 is even more extreme, with the equilibrium population close to the optimum for Environment 1 and poorly adapted to Environment 0.

Transmission-selection covariance can also shift a population from one “adaptive peak” to another. Figure 5 shows a case in which expected fitness exhibits two distinct peaks, one higher than the other (*c* = 4, *a* = .6 in Equations 12). In the example, expected fitness is maximized with *ϕ* = 0 in Environment 0, but the greater heritability in Environment 1 means that the population adapts to that environment, even though half of the population ends up in Environment 0 and and has nonzero heritability there.

**Figure 5:**
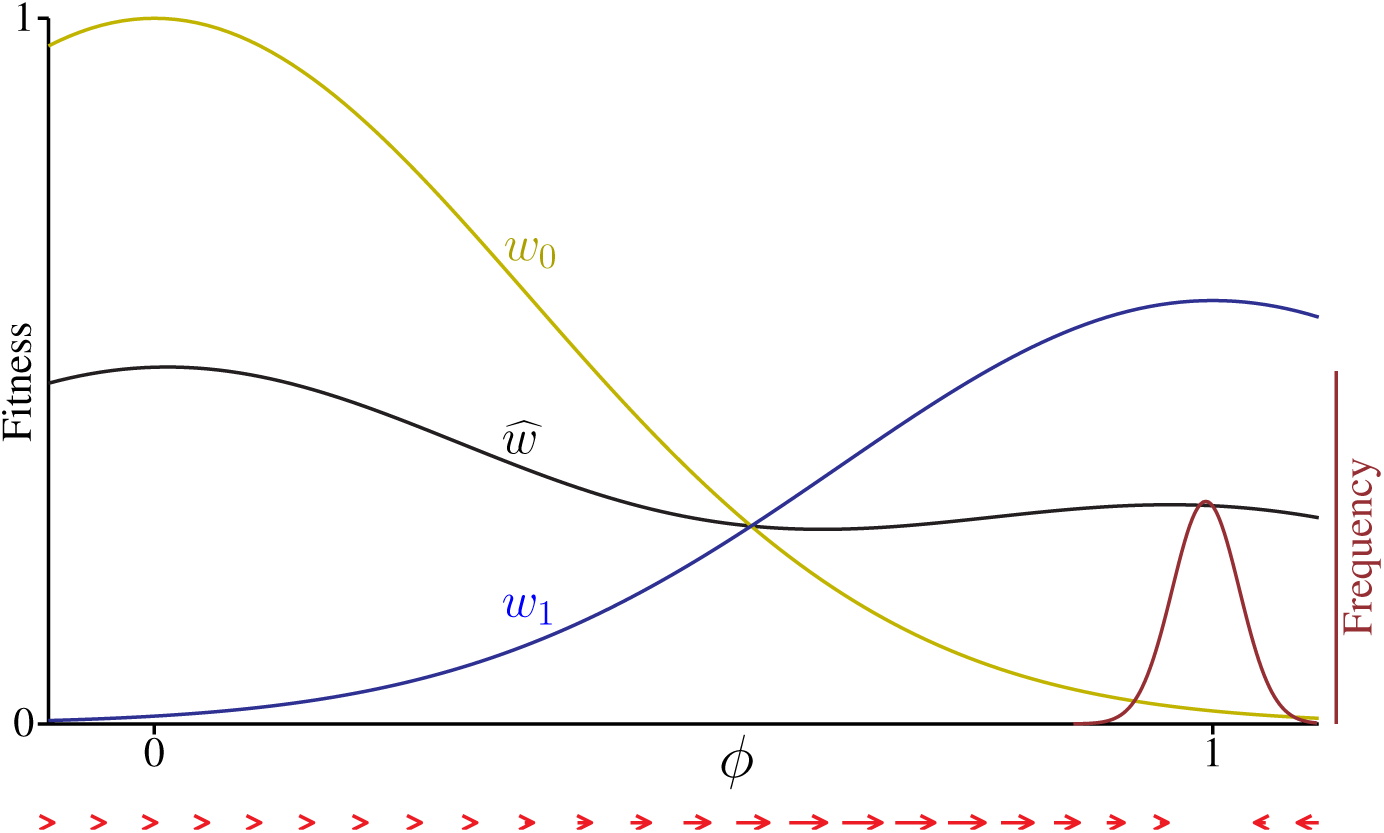
Different heritabilities with multiple peaks. Fitness in Environment 0 (*w*_0_), in Environment 1 (*w*_1_), and the expected fitness 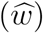 for an individual that has a probability of 0.5 of finding itself in either environment. The distribution (purple) is the equilibrium distribution when 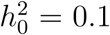 and 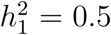. The arrows show the magnitude and direction of 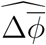

All of the heritability values used in the preceding examples are well within the range of values observed in natural populations.

In the examples in Figures 4 and 5, we held the heritability within each environment fixed so as to highlight the consequences of variation between environments. In fact, heritability and additive genetic variance can themselves evolve. In particular, it is often assumed that heritability and additive genetic variance will decline under stabilizing selection.

To see the evolutionary consequences of change in transmission, note that the positions of the equilibrium distributions are determined by the balance of two different “forces”, captured by the two terms on the righthand side of Equations 11. The first term, 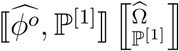 (or, equivalently, 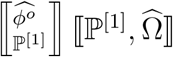 involves only the expected values of “additive genetic variance” (or heritability) and fitness. Decreasing the expected additive variance decreases the absolute value of this term, making selection less effective. By contrast, the second term,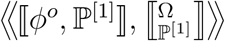 (equivalently 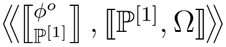) is influenced by the variation in additive variance, but not by the mean value.

The consequences of reducing additive variance will thus depend on whether the absolute values in each environment, or the difference between them, declines faster. In the simple case in which additive variance declines by the same proportion in each environment, the two terms on the righthand sides of Equations 11 will decline to the same degree, so the equilibrium will be unchanged.

The example in Figure 4 shows the effects of changing the variance in heritability while holding the mean constant. For comparison, Figure 6 illustrates the effects of changing only the mean heritability. Increasing the mean of *h*^2^ shifts the equilibrium distribution back towards what would be predicted if *h*^2^ were constant. Note, though, that even with a relatively high mean heritability, moderate variation leads to the population being better adapted to Environment 1.

**Figure 6:**
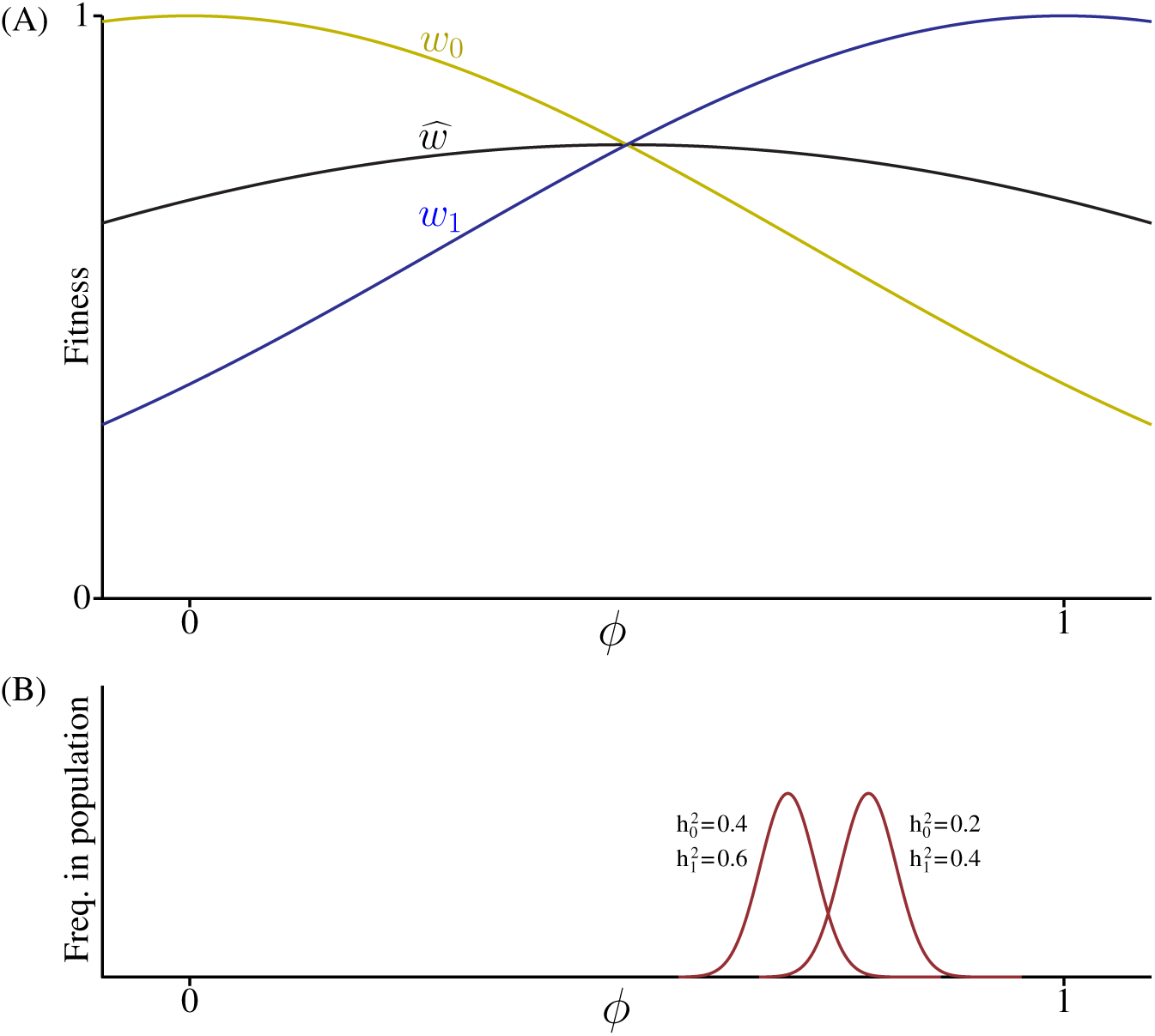
Effects of changing mean heritability. (A): Fitness in Environment 0 (*w*_0_), in Environment 1 (*w*_1_), and the expected fitness 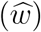. As in Figure 4A. (B): Equilibrium phenotype distributions for two cases with the same value of 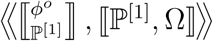 (since the difference between 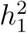 and 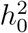 are the same in each case), but different values of 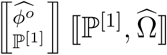.

**Figure 7:**
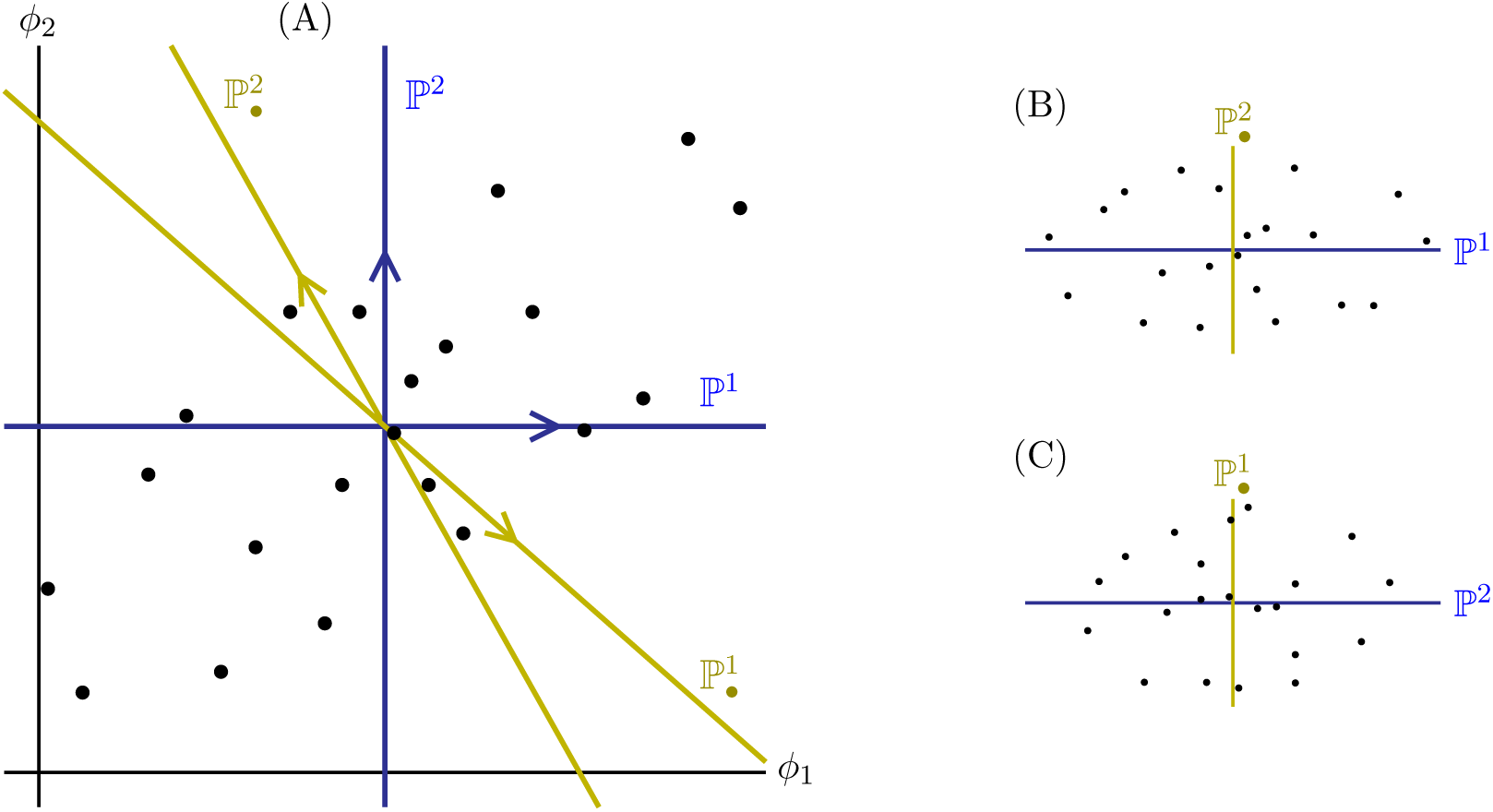
Biorthogonal bases: (A) A joint distribution of values for two traits, *ϕ*_1_ and *ϕ*_2_, and the corresponding first order simple (blue) and conditional (yellow) bases. Though ℙ^1^ and ℙ^2^ are at right angles, they are not orthogonal with respect to this distribution of points. (B) The same points projected onto one simple base (ℙ^1^) and the orthogonal conditional base 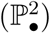. The points are uncorrelated in this space because these bases are orthogonal with respect to these points. (C) The distribution of points projected into ℙ^2^ and 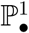. The points are again uncorrelated in this space.

In the examples given above, individuals have a fixed probability 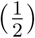 of encountering each environment. We might expect that once individuals become better adapted to one environment, there would be selection on other traits to increase the probability of encountering that environment. In order to investigate this, we need to generalize Equations 9 to apply to multiple traits.

### Joint evolution of multiple traits

Multivariate orthogonal polynomials pose a new challenge. Though they are relatively easy to find, constructing an orthogonal basis requires choosing a way to order our traits, and the ordering that we choose influences what terms show up in our final result. For example, if we choose the order (*ϕ*_1_, *ϕ*_2_, *ϕ*_3_) we will have a term for the effects of *ϕ*_2_ independent of *ϕ*_1_ but not independent of *ϕ*_3_.

The way around this problem is to find a pair of bases with the property that, for each basis, each axis in it is orthogonal to all but one of the axes in the other basis. A pair of bases that have this property are said to be “biorthogonal” to one another (Dunkl and Xu, 2001), and it is relatively easy to construct a pair of biorthogonal bases for our phenotype space (see Supplementary Material). Figure 18 shows a simple example of a pair of bases that are biorthogonal to one another. Given such a pair of bases, we can find the covariance between two variables, such as *ϕ°* and Ω, by projecting one variable into one basis, and the other variable into the biorthogonal basis.

For multivariate polynomials, we modify our notation a bit. The superscript (without brackets) denotes the trait, and the number of superscripts shows the degree of the polynomial. So ℙ^*i*^ is a first order polynomial containing trait *ϕ*_*i*_, and ℙ^*ij*^ is a second order polynomial containing *ϕ*_*i*_*ϕ*_*j*_.

Table 2 shows the pair of biorthogonal bases that we use (see Methods). The first order “simple” bases are just the mean centered trait values, while the first order “conditional” bases are constructed so that 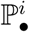 is orthogonal to all of the simple bases except ℙ^*i*^. ℙ^1^ and ℙ^2^ are not orthogonal to one another, nor are 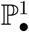 and 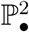, but the conditional basis 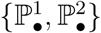 is biorthogonal to the simple basis {ℙ^1^, ℙ^2^}. The simple and conditional second order polynomials have a similar relationship to one another (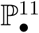 is orthogonal to ℙ^12^ and ℙ^22^), and in addition all second order polynomials are orthogonal to all first order polynomials.

**Table 2:**
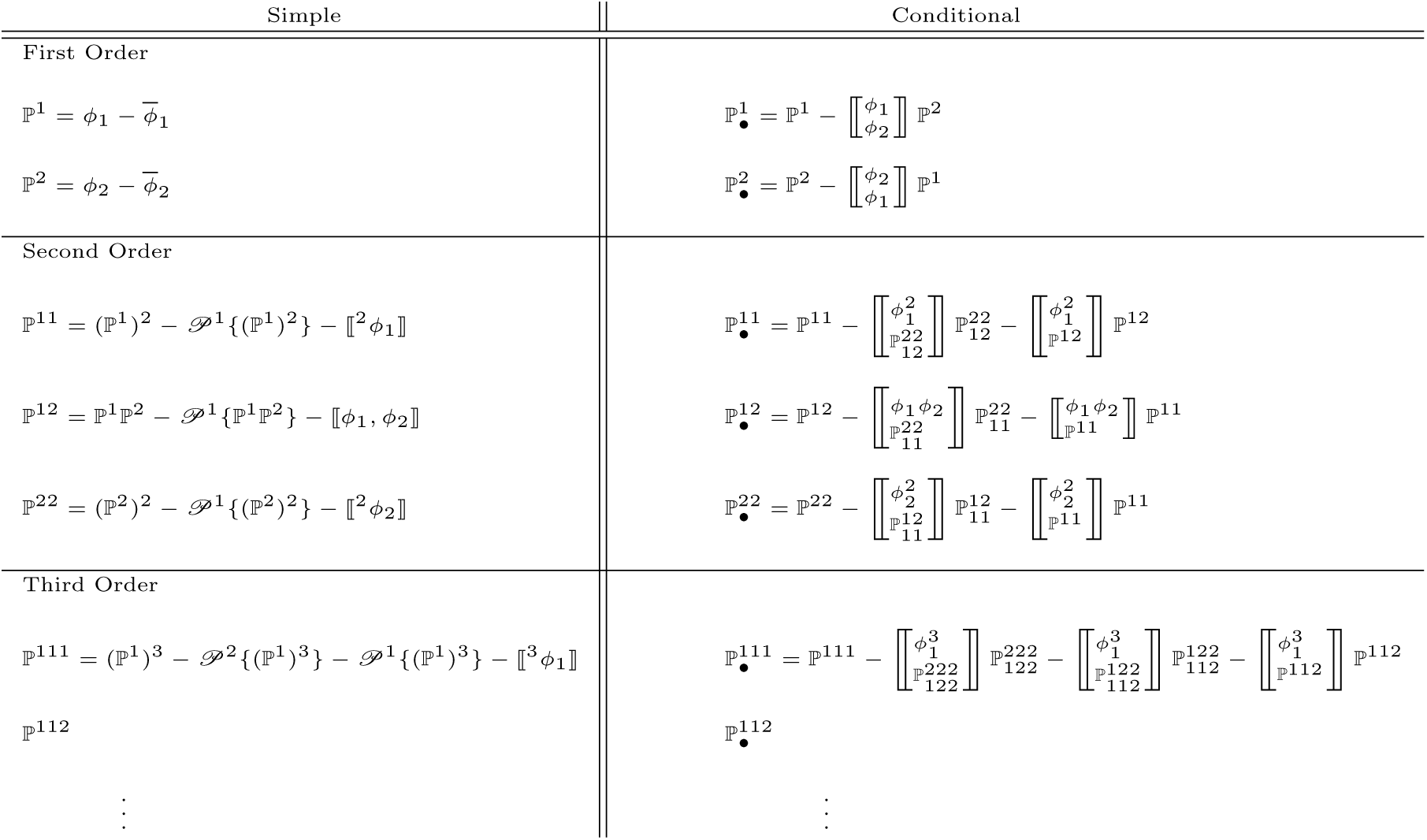
The polynomials on the left (simple) form a non-orthogonal basis. The polynomials on the right (conditional) form a different non-orthogonal basis that is biorthogonal to the basis defined by the polynomials on the left. 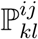 is the polynomial with first term *ϕ*_*i*_*ϕ*_*j*_ that is orthogonal to *ϕ*_*k*_*ϕ*_*l*_.

The general rule is: 1) The polynomials within each order are orthogonal to all those of other orders, and 2) within each order, the polynomials on the right and left are biorthogonal.

Regressing a variable, such as 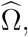, on the simple basis ℙ^1^ yields the simple regression of 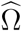 on *ϕ*_1_. By contrast, regressing on the conditional basis 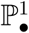 yields the *partial* regression of 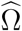 on *ϕ*_1_ (Saville and Wood, 1991). Regressing on 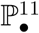 is the equivalent of taking the partial regression on 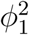 if we were to include all first and second order combinations of *ϕ*_1_ and *ϕ*_2_ in the analysis.

In this way, we can build up two bases that are biorthogonal to one another and for which the regression terms have straightforward biological interpretations. (A note on terminology: Our “simple” and “conditional” bases are analogous to the “contravariant” and “covariant” bases used in tensor analysis in physics (Dodson and Poston, 1991). They are not exactly the same, though, since we are not normalizing the inner products).

As noted above, the reason for constructing biorthogonal bases is to be able to write a general equation for the covariance between fitness and offspring phenotype. If we write offspring phenotype in terms of the simple basis, and relative fitness in the conditional basis, then for a set of *m* traits we find the vector of expected changes in all trait means to be (see Methods):

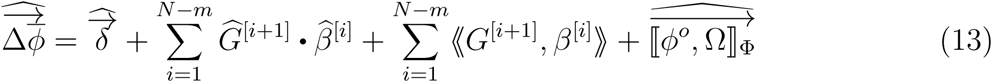

(Where 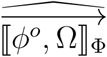 is a vector, the *j*^*th*^ element of which captures the average covariance between 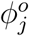 and Ω within sets of individuals that share the same value of every phenotypic trait).

Here, *G* is a tensor containing the covariances between offspring and parents, and *β* a tensor containing the regressions of relative fitness on the traits. The notation *a* ·*b* denotes the inner product of two tensors (see Methods). If *a* is a matrix and *b* a vector, then the inner product is the same as standard pre-multiplication of the vector by the matrix. (For our purposes, a “tensor of degree *n*” is an *n* dimensional array of values. A tensor of degree 1 is a vector, and a tensor of degree 2 is a square matrix.) Specifically:

*G*^[*n*]^: This is a tensor of degree *n* (so *G*^[2]^ is a matrix) that contains all of the covariances of offspring phenotype with all of the (*n* −1)^*st*^ order *simple* polynomials. *G*^[2]^ is just the offspring-parent covariance matrix, which is essentially the same as the standard *G* matrix used in quantitative genetics (though note that *G*^[2]^ need not be symmetrical, since in general 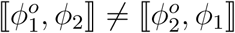 (Rice, 2004))

For the case of two traits, *ϕ*_1_ and *ϕ*_2_, *G*^[2]^ is:

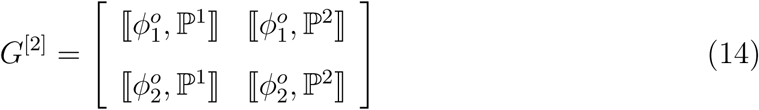

Nonlinear relationships between offspring and parents are captured by the higher degree *G* tensors. For example second order effects of parents on offspring are captured by the elements of *G*^[3]^, with 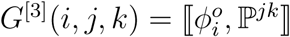.

*β*^[*n*]^: A tensor of degree *n* (so *β*^[1]^ is a vector) containing the regressions of relative fitness (Ω) on the *n*^*th*^ order *conditional* polynomials. *β*^[1]^ is just the fitness gradient:

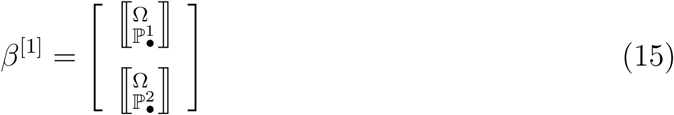

Higher order *β* tensors contain nonlinear effects of phenotype on relative fitness. For example, *β*^[2]^ is the matrix:

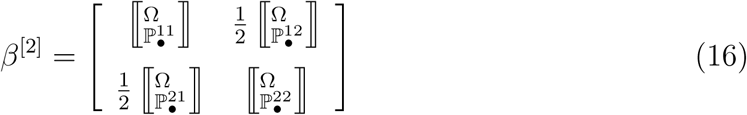

The “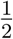” terms in the matrix in Equation 16 account for the fact that 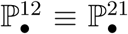 and that this term appears only once in each summation in Equation 13. For higher order *β* tensors, we similarly divide each element by the number of times that equivalent elements appear in the array. In general, for *m* different traits, the elements of *β*^[*n*]^ are given by:

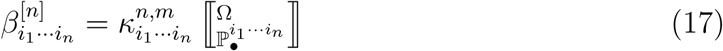

To calculate the combinatorial term, *κ*, we denote by *x*_*i*_ the number of instances of index *i* among the subscripts (*e.g* for 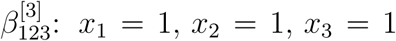, and for 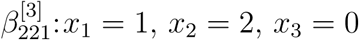). If we are considering *m* different traits, then the combinatorial term *κ* is:

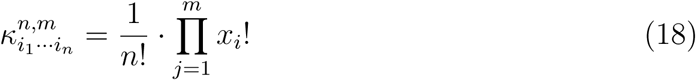

(keeping in mind that 0! = 1).

If we are considering only one trait, then *m* = 1 in Equation 13 and it reduces to the univariate Equation 9.

Because we projected relative fitness into the conditional basis, we end up with regressions of relative fitness on the ℙ_•_ polynomials. As noted, these are equivalent to partial regressions (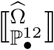 is what we would get for the partial regression of 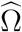 on *ϕ*_1_*ϕ*_2_ if we included only first and second order terms in our analysis). These terms capture the shape of the relative fitness “landscape”.

(Note that we could, just as easily, represent fitness in the simple basis and offspring phenotype in the conditional basis. This would capture selection with covariances (the first order term being the “selection differential”) and transmission with partial regressions of offspring on parents. Note that these would not be heritabilities in the conventional sense (in which heritability is a simple, not partial, regression), but could yield insight by allowing us to distinguish the independent contributions of different traits to transmission. In particular, measuring offspring phenotype in the conditional basis would allow us to identify traits that influence the transmission of other traits.)

Equation 13 allows us to work with any fitness distribution and any transmission rules. The variables *ϕ, ϕ°*, and Ω may be continuous or discrete, and *ϕ°* and Ω may be random variables, having distributions of possible values. Below, we illustrate application of this result with an extension of the variable heritability case discussed earlier.

### Unequal heritabilities combined with habitat selection

We return to the case of a trait that exhibits different heritabilities in different environments, but now add a second trait that influences the probability that an individual finds itself in a particular environment. Heritability can now change as this “habitat preference” trait evolves.

We consider a trait, *ϕ*_*s*_, that influences fitness differently in the two habitats, with *ϕ*_*s*_ = 0 having the highest fitness in habitat 0, and *ϕ*_*s*_ = 1 having the highest fitness in habitat 1. Specifically, fitness in habitat 0 (*w*_0_) and in habitat 1 (*w*_1_) are given by:

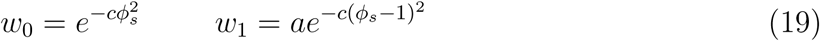

As before, we consider a case in which inheritance is linear within each environment, with 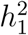 being the heritability in environment 1, and 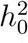 being the heritability in environment 0. Now, however, the trait *ϕ*_*h*_ determines the probability that an individual ends up in Environment 1. This might be a behavioral trait such as habitat preference, or a physiological trait that influences the probability of germinating or hatching in different habitats. Formally, we write the probability of encountering environment 1 as:

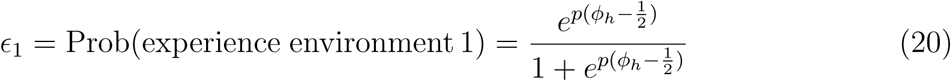

Where the parameter *p* determines how abruptly habitat preference changes as *ϕ*_*h*_ changes (Figure 8).

Relative fitness, 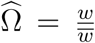, is now determined by *ϕ*_*s*_, *ϕ*_*h*_, and *ϵ*_1_. Denoting the relative fitness of an individual in environment *i* as Ω_(*i*)_, we have:

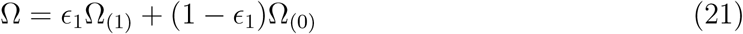

**Figure 8:**
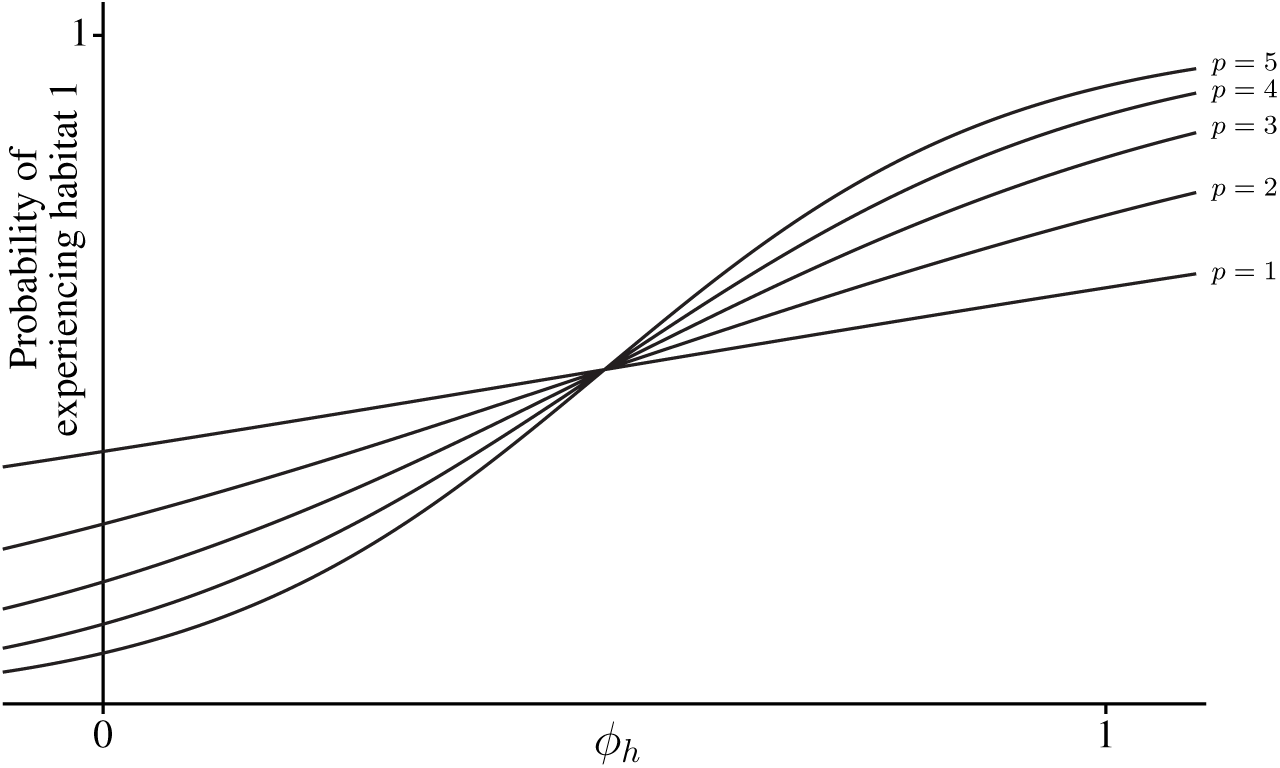
Habitat preference: Probability of using habitat 1 as a function of the trait *ϕ*_*h*_, for different values of *p* in Equation 20

This produces a fitness surface with two peaks, one at (*ϕ*_*s*_ = 0, *ϕ*_*h*_ = *-∞*) and one at (*ϕ*_*s*_ = 1, *ϕ*_*h*_ = *∞*)). Figure 9 shows the surfaces for two different values of *p*. Note that increasing the effect of *ϕ*_*h*_ on habitat selectivity deepens the valley between the two adapted states.

Because heritability may be different in the different habitats, the value of trait *ϕ*_*s*_ among an individual’s offspring 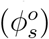 is a function of both *ϕ*_*s*_ and *ϕ*_*h*_ in the parents. We can write the mean values of *ϕ*_*s*_ and *ϕ*_*h*_ among an individual’s offspring as:

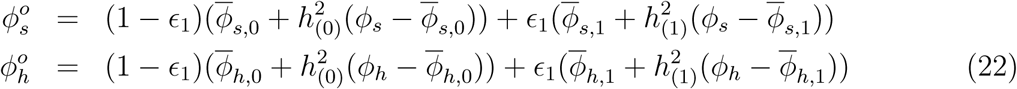

where 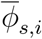 is the mean value of trait *ϕ*_*s*_ in environment *i*.

**Figure 9:**
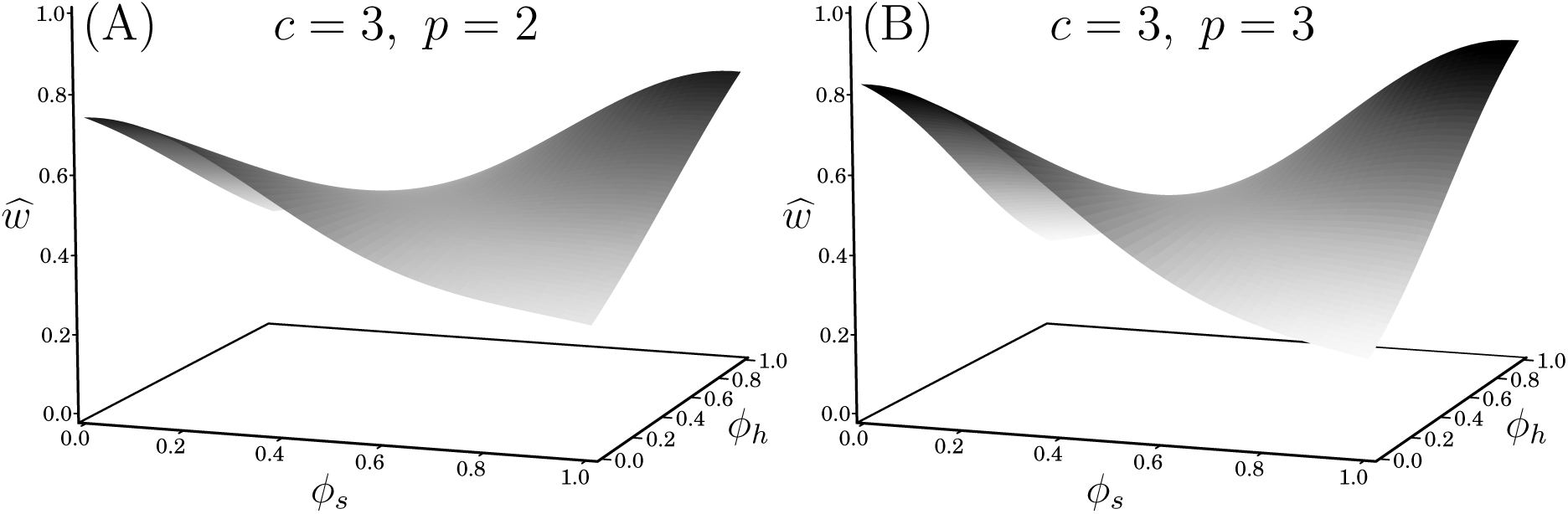
Expected fitness as a functions of *ϕ*_*s*_ and *ϕ*_*h*_: Fitness surfaces for the case in which *c* = 3 in Equation 19 for two different habitat selection functions (values of *p* in Equation 20). In (A) *p* = 2. In (B), *p* = 3.

Equations 22 show that 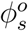 is a nonlinear function of *ϕ*_*s*_ and *ϕ*_*h*_ (since *ϵ*_1_ is a function of *ϕ*_*h*_). We thus expect that, in addition to the terms in Equation 11, there will be a second order term capturing the joint effects of *ϕ*_*s*_ and *ϕ*_*h*_, namely 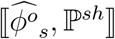. We also expect 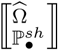 to be nonzero, since fitness depends – in a nonlinear way – on both *ϕ*_*s*_ (the trait under different selection in the different environments) and on the environment encountered (which is influenced by *ϕ*_*h*_). If, for simplicity, we consider the case in which the traits *ϕ*_*s*_ and *ϕ*_*h*_ are normally distributed and uncorrelated, then Equation 13 simplifies to:

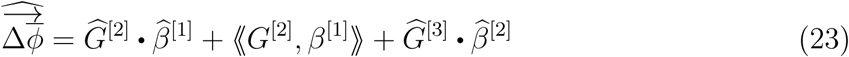

(There is no ⟪*G*^[3]^, *β*^[2]^⟫ term because, in a large population, there will be no appreciable stochastic variation in *β*^[2]^).

Figure 10 shows the vector fields for evolutionary change, on the fitness landscape shown in Figure 9, for the case in which heritability is the same in both environments and a case in which heritability is larger in environment 1. (Note that both traits could continue to change beyond the range shown).

**Figure 10:**
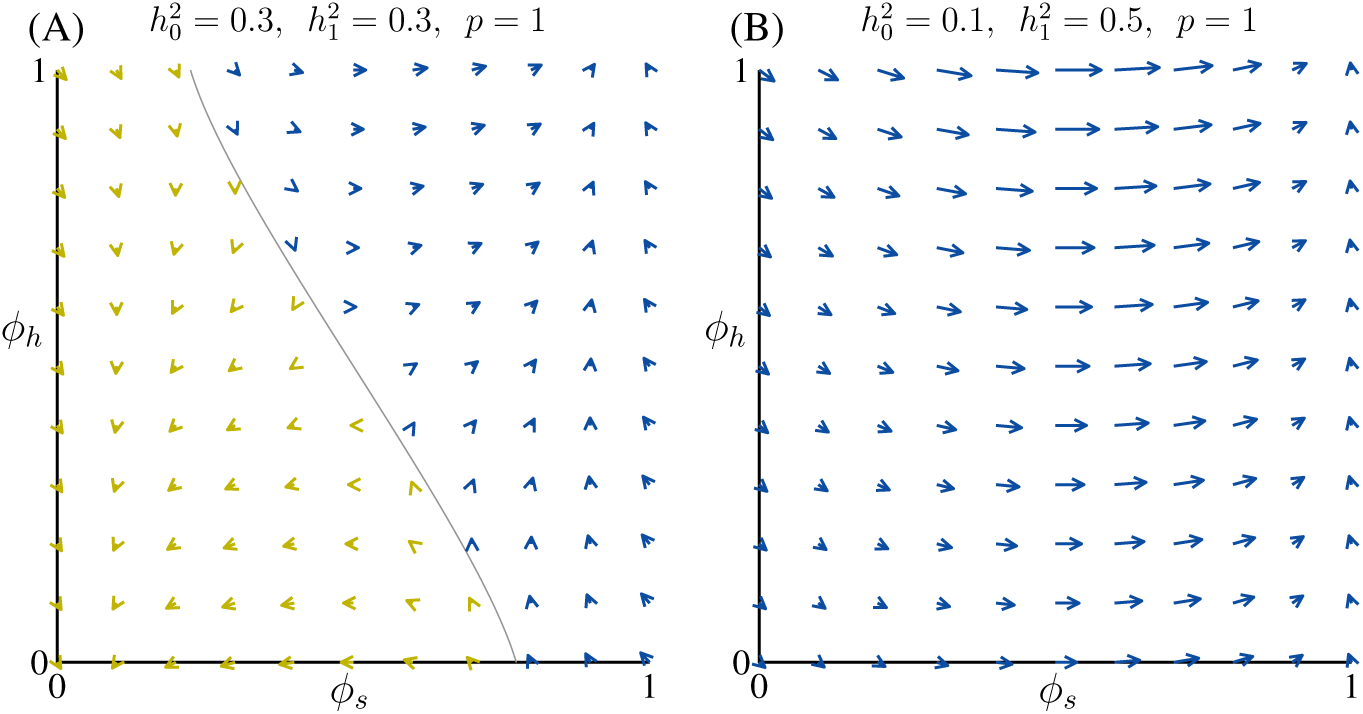
Joint evolution of a selected trait and habitat preference: Vectors showing the direction of evolution for a case in which *c* = 3 in Equation 19 and habitat preference is relatively weak (*p* = 1 in Equation 20 and Figure 8). The vectors are magnified by a factor of 10. In (A), heritability is the same in both environments. In (B), heritability is higher in environment 1 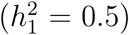 than in environment 0 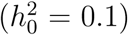. Yellow vectors indicate points from which the population will evolve to a point at which it is well adapted to, and prefers to use, habitat 0. Blue vectors indicate points from which the population will evolve to be well adapted to, and prefer, environment 1.

When heritability is the same in the two environments (Figure 10A), then the system behaves like a classic case of evolution on a landscape with two peaks – the population always evolving “uphill” on the fitness surface. By contrast, when heritability is higher in Environment 1 (Figure 10B), all initial conditions in the range shown ultimately evolve towards adaptation to, and preference for, environment 1. As in the univariate case, a correlation between transmission and selection can pull a population across an adaptive “valley”. Here, though, habitat preference can evolve as well, making it possible for the population to evolve to use the higher heritability environment exclusively.

There is no selection for increasing heritability *per se*. Differential heritability causes the population to become better adapted to the environment with higher *h*^2^, and then selection favors increasing use of the habitat to which individuals are better adapted. Heritability itself evolves as a byproduct.

Interestingly, increasing the intensity of habitat selection – by increasing the value of *p* in Equation 20 and Figure 8 – reduces the range of initial conditions from which the population is expected to evolve towards *ϕ*_*s*_ = 1, *ϕ*_*h*_ = 1 (Figure 11). The reason is that, with strong habitat selection, individuals that initially prefer habitat 0 rarely encounter habitat 1, so there is little opportunity to adapt to it – regardless of the heritability values.

**Figure 11:**
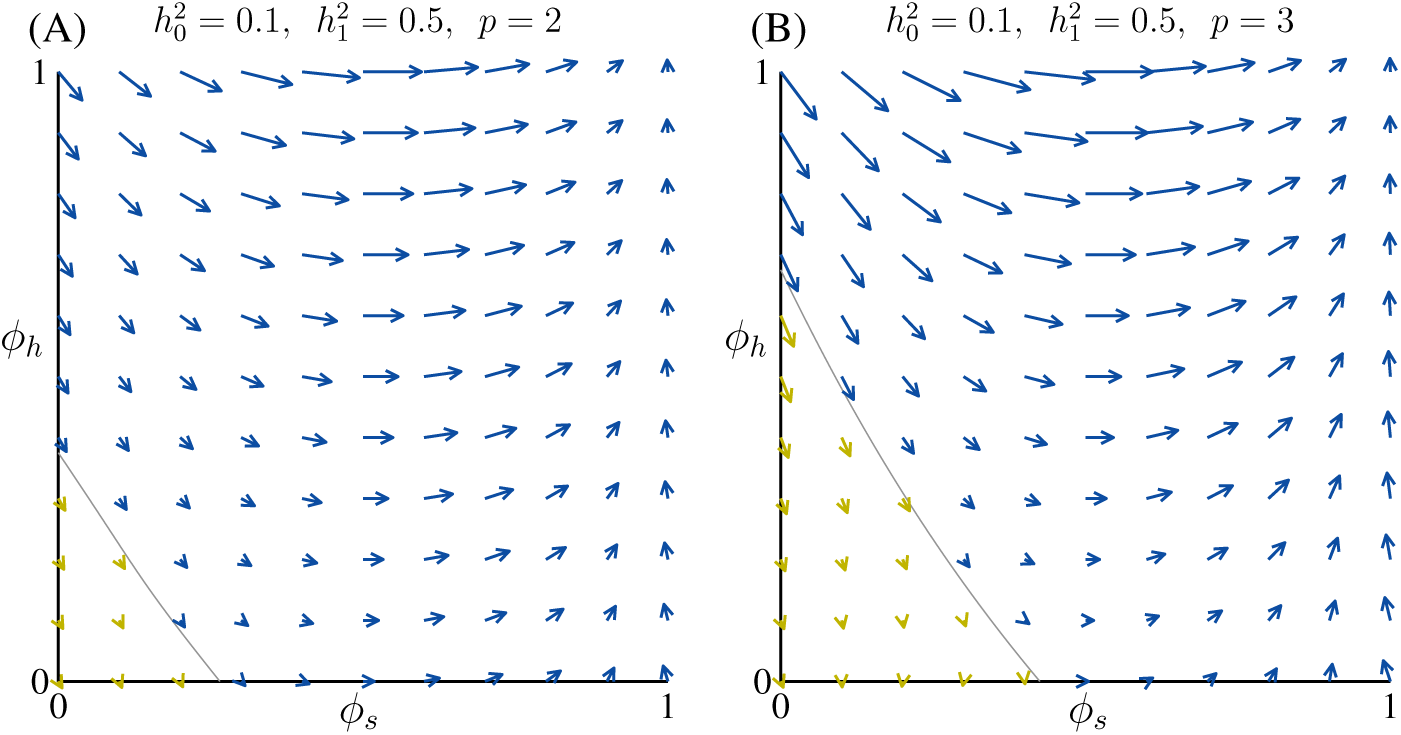
Effects of changing strength of habitat preference: (A) Same as in Figure 10(B) except with stronger habitat selection (*p* = 2 instead of 1). (B) Same as in Figure 10(B) except with *p* = 3.

Figure 12 shows the contributions of linear deterministic, stochastic, and nonlinear deterministic components (the three terms on the righthand side of Equation 23) to the total change for the case shown in Figure 10B. The vector field in Figure 12(A) shows the contribution of the term *G*^[2]^ • *β*^[1]^ from Equation 13. This is what a standard multivariate quantitative genetics model would yield for this system. This term pushes a population up-hill on the fitness surface, though at different rates for different values of *ϕ*_*h*_, which influences the additive genetic variance.

**Figure 12:**
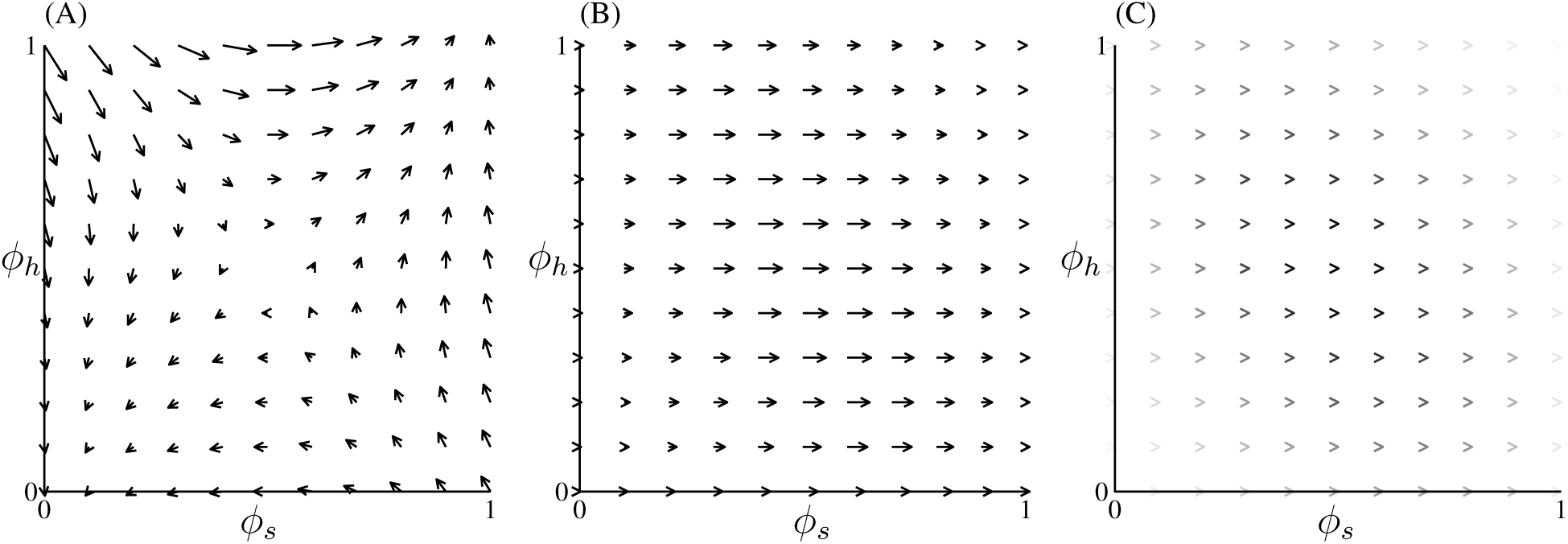
Components of the vector field in Figure 11B: (A) shows the vector field resulting from only considering terms like 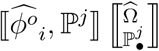. This is what a deterministic quantitative genetics model would predict. The vectors are not exactly symmetrical because the “additive genetic variance” varies across the range of values of *ϕ*_*h*_. (B) shows the contribution of terms like 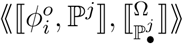. (C) shows the contribution of second order terms like 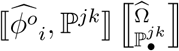. These are much smaller than the vectors in (A) and (B), so magnitude is represented by shading, with darker arrows being larger.

The contribution of ⟪*G*^[2]^, *β*^[1]^ ⟫ – the probability covariance between additive genetic variance and selection – is shown in Figure 12(B). Note that the maximum magnitude of these vectors is similar to that of the direct selection vectors in Figure 12(A). Furthermore, this vector field is of greatest magnitude in the vicinity of the “saddle” of the fitness surface, where direct selection is relatively weak. (These vectors have no component in the *ϕ*_*h*_ direction because, within each environment, there is no selection on *ϕ*_*h*_ – it influences fitness only through its effect on the probability of experiencing one or the other environment.)

The second order component of transmission, captured by G^[3]^ • *β*^[2]^ in Equation 13, while nonzero, contributes relatively little to overall change in this particular system (Figure 12(C)). Notably, though, the contribution of second order terms – though small – is also maximal in the region in which directional selection is weak.

In the examples discussed above, there is no correlation between the traits *ϕ*_*s*_ and *ϕ*_*h*_. In reality, joint selection on different traits is expected to alter the covariance between them (Steppan et al., 2002; Arnold et al., 2008; Melo and Marroig, 2015). This is apparent when we generalize Equation 8 to give the expected change in the covariance between the two traits:

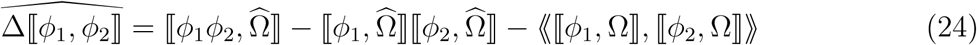

The first term on the righthand side of Equation 24 captures selection on the product of the two traits. The second term captures the product of independent selection on each trait. Note that this second term contributes negatively; meaning that independent selection to increase both traits, or to decrease both traits, leads to a negative covariance between them. This is a generalization of the Hill-Robertson effect, whereby independent selection for alleles at two different loci creates negative gametic disequilibrium between them. The third term captures the probability covariance between selection on the two traits.

For the system considered here, Equation 24 is positive for all values of *ϕ*_*s*_ and *ϕ*_*h*_ in the range [0, 1]. (This is not surprising, since the fitness surface (Figure 9) is saddle shaped). Introducing such a correlation, in this case, causes the 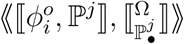 vector field and the 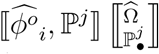 vector field to cancel one another out more in the region of high *ϕ*_*h*_ and low *ϕ*_*s*_ (upper left in the figures). This reduces somewhat the range of initial values from which a population is expected to evolve toward use of habitat 1 (Figure 13A).

**Figure 13:**
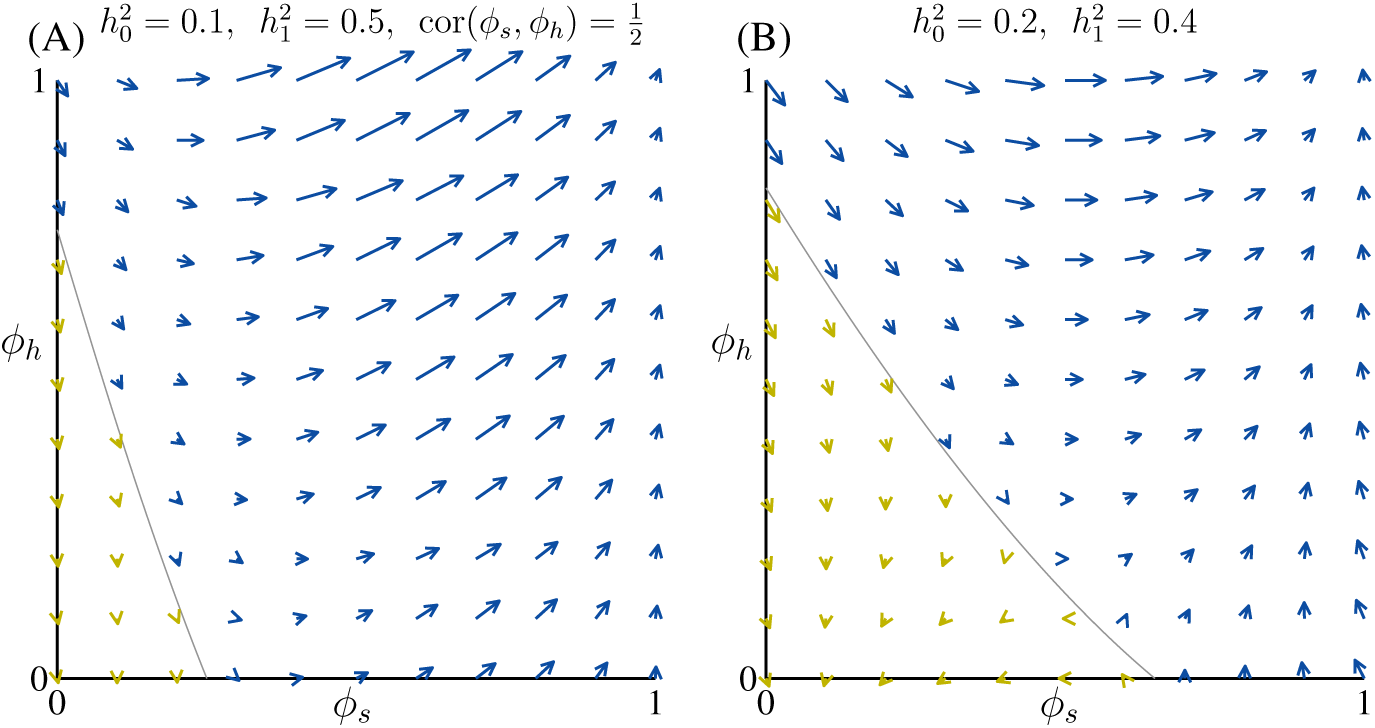
Effects of correlation between *ϕ*_*h*_ and *ϕ*_*s*_, and changing the variance in heritability: (A) Case in which *ϕ*_*h*_ and *ϕ*_*s*_ are positively correlated. All other parameters are as in Figure 11(A). (B) As in Figure 11(A) but with lower variance in heritability, here, 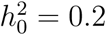 and 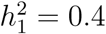.

The magnitude of the transmission selection covariance increases with both the variance in selection and the variance in transmission. Thus, reducing the difference in heritability between the environments reduces the size of the region from which populations tend to evolve towards *ϕ*_*s*_ = 1, *ϕ*_*h*_ = 1 (Figure 13B).

## Discussion

The method of projection onto orthogonal or biorthogonal polynomials can be applied to any kind of variable – continuous or discrete, deterministic or stochastic (Jackson, 1941). If these polynomials are themselves defined by the distribution of variation within a population, then they form the basis of a natural space within which to study population level processes.

Projecting relative fitness and offspring phenotype into this space allows us to apply general results, such as the stochastic Price equation (Equation 1), to multivariate cases with complex selection and transmission. Because the space is constructed from the population distribution itself, we need not make simplifying assumptions about normality, linearity, or continuity.

The univariate (Equations 10 and 9) and multivariate (Equation 13) results that we derive allow for any patterns of selection or transmission. General results of this sort can yield insights into the mechanics of evolution – such as by revealing new evolutionary processes – but are unwieldy for studying particular cases. By introducing simplifying assumptions after deriving the main result, though, a general theory of this sort can serve as a formula for constructing special case models. In this paper, we use our general result to derive models of evolution in which both selection and transmission are influenced by the environment in which organisms find themselves.

Various processes may lead to similar populations exhibiting different heritabilities in different environments. Likely causes include differences in environmental variance (Lynch and Walsh, 1998), and nonlinear effects of environment on phenotype (Holloway et al., 1990; Rice, 2012). Whatever the cause, a consequence is that the response to selection will vary with environment.

By itself, the phenomenon of different heritabilities in different environments produces no directional evolutionary change. A directional effect emerges only when heritability (or additive genetic variance) covaries with selection. Such a transmission-selection covariance is likely to be common, though, since both are often influenced by variable environmental factors.

When transmission and selection do covary, the effect can be pronounced, with modest differences in heritability leading the population to become much better adapted to those environments in which heritability is highest. One consequence of this is that the evolutionary equilibrium that a population approaches can be quite different from that predicted by models, such as evolutionary game theory, that treat heritability as fixed.

When habitat use is itself allowed to evolve, modest differences in heritability between environments can lead to organisms exclusively using environments with higher heritability, since adaptation to such environments leads to selection for any other traits that increase individuals’ use of those environments. This gives insight into one mechanism through which heritability itself evolves – as a byproduct of selection on habitat choice. These results also suggest one explanation for the observation that stressful environments often confer low heritability (Charmantier and Garant, 2005; Husby et al., 2011). Adaptation to environments that facilitate higher heritability will lead to other environments being “stressful” simply because organisms are poorly adapted to them.

The interplay of stochastic transmission and selection described here is quite different from the “directional stochastic effects” that result from stochastic variation in fitness alone (R<u>ic</u>e, 2008; Rice et al., 2011). In particular, because heritability (unlike 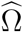) is not a function of 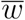, the effects of variable heritability are independent of population size.

### Axiomatic theories and model building

All models of evolution make assumptions about transmission. In some cases, such as classical population genetics, these assumptions are so entwined with the model of selection that it is not obvious from the final result that a very specific transmission process (such as Mendelian transmission with random mating) is assumed. In other cases, such as much of quantitative genetics theory, transmission is explicitly represented – but is captured only by a single parameter, heritability or additive genetic variance, that is then often treated as a constant. Both of these approaches can obscure the role that transmission plays in shaping the dynamics of evolution.

We have taken a somewhat different approach. By using a very general method for capturing relationships between variables, we can represent both offspring phenotype and parental fitness in ways that allow any transmission pattern to be combined with any selection regime. The result is essentially a multivariate stochastic version of the exact breeder’s equation (Heywood, 2005), and it allows for a more complete account of how transmission and selection jointly determine the path of evolution.

## Methods

### Orthogonal polynomials

Orthogonal polynomials are constructed so that each is independent of (orthogonal to) all of the others. For functions, orthogonality is defined relative to a “weight function”. There are a number of sets of classical orthogonal polynomials, each defined with respect to a different weight function. For example, the Legendre polynomials used to study function valued traits such as continuous growth processes (Kirkpatrick et al., 1990; Pletcher and Geyer, 1999; Meyer and Kirkpatrick, 2005; Griswold et al., 2008; Stinchcombe et al., 2012) have as a weight function a uniform distribution on a fixed interval. The key to the approach presented here is to use the population itself as the weight function (see the Supplementary Material).

For a phenotypic trait *ϕ*, the *n*^*th*^ order polynomial is a function containing *ϕ*^*n*^ and lower powers of *ϕ*, constructed so that it is orthogonal to all lower order polynomials. Denoting the *n*^*th*^ central moment of the population distribution by ⟦^*n*^*ϕ*⟧, and the simple regression of *a* on *b* as 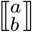, the orthogonal polynomials corresponding to different powers of *ϕ* are given by:

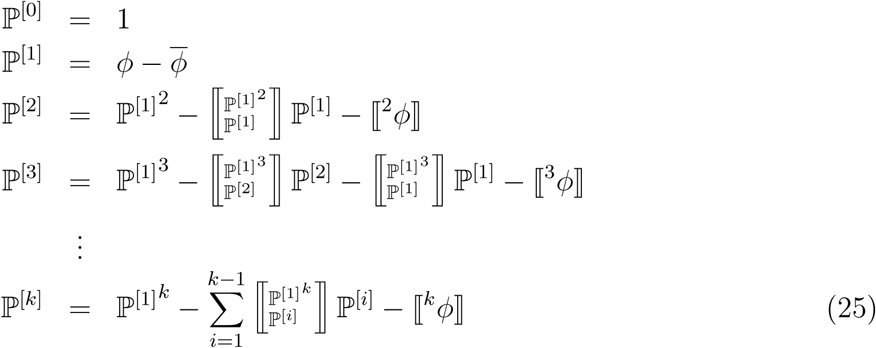

Equations 25 show how higher order orthogonal polynomials can be derived sequentially from lower order ones, using the Gram-Schmidt process (see Supplementary Material).

### Rules for combining frequency and probability operations

Frequency operations, such as 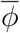 and ⟦^2^*ϕ*⟧ (the mean and variance of phenotypes currently in the population) are calculated over a set of individuals. Probability operations, such as 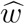 and ⟪^2^*w* ⟫ (the mean and variance of fitness), are calculated over the probability distribution of possible values. Both kinds of operations follow the same rules relating means, variances, and covariances. For example:

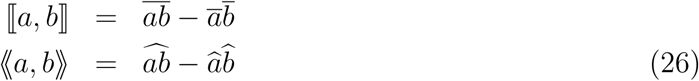

Some variables, such as offspring phenotype and fitness, are both frequency and probability variables. In such cases, we need to combine frequency and probability operations. In the following derivations, we will make use of two fundamental rules (Rice and Papadopoulos, 2009).

First, the order of frequency and probability means can be reversed:

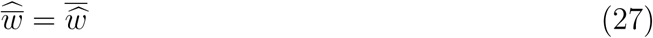

Second, for two random variables, *a* and *b*, frequency and probability covariances are related by the following rule:

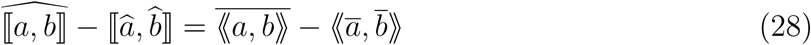

If *c* is a non stochastic variable (such as phenotype of parents, *ϕ*, which is a fixed value for an individual but varies across a population), then:

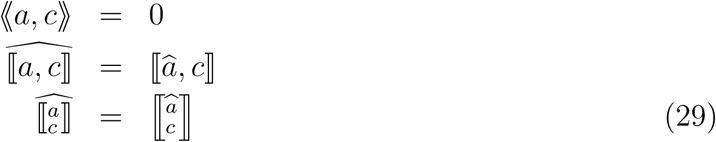

### The core equation

All of our main results follow from a fundamental relationship between offspring phenotype, relative fitness, and and the mean phenotype in the next generation. In the absence of immigration, the expected value of the mean of a trait *ϕ* after one generation 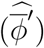 is given by (Rice, 2008):

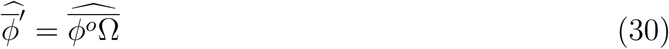

Where 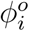 is the mean phenotype of individual *i*’s offspring, and 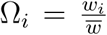 (conditional on 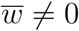) is the relative fitness of individual *i*.

### Selection differential for the variance

With no loss of generality we can shift our axes so that 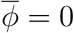, which means that 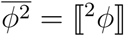. If we are concerned only with the selection differential (the change due only to selection, prior to reproduction) then *ϕ*^*o*^ = *ϕ*. To find the selection differential for the variance, note that the expected phenotypic variance after selection, 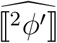, is given by:

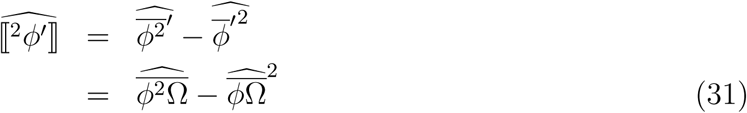

Note that we are using the fact that *ϕ*^2^ can be thought of as just another trait. Now use the facts that 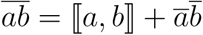, and that 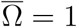:

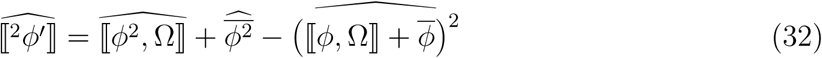

Since we set 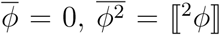, the initial phenotypic variance. Subtracting this ⟦^2^*ϕ*⟧ from both sides yields:

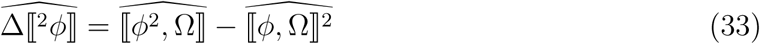

Finally, we apply the Rule 29 (since *ϕ* is not a stochastic variable) to the righthand side of Equation 33,

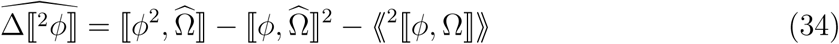

### Single phenotypic trait

We need to write Equation 30 in terms of *ϕ*.

For individual (or mated pair) *i*, we write 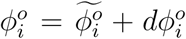, and 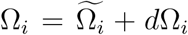. Where *dx*_*i*_ is the difference between *x*’s value of *i* and the mean of individuals that share the same phenotypic value (*ϕ*) as does *i*. Because 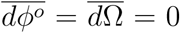, and the *dx* values are independent of the 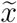 values, we get:

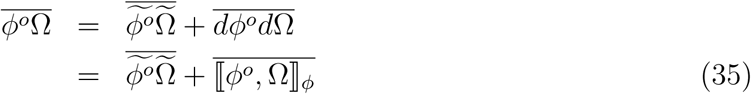

Where 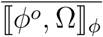 is the average, across all phenotypes, of the frequency covariance between *ϕ*^*o*^ and Ω for individuals who share the same value of *ϕ*, weighted by the number of individuals with that value.

Using Equation 3 for 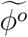 and Equation 7 for 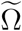, we find:

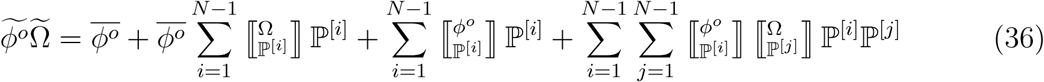

(See Supplimentary Material)

Now recall that the ℙ terms were constructed so as to be orthogonal to one another and to each have a mean of 0 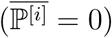. This means that

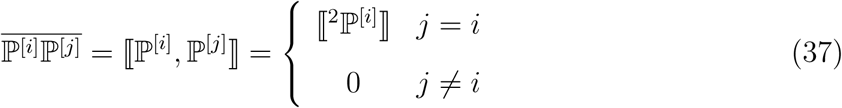

The frequency mean of Equation 36 thus reduces to:

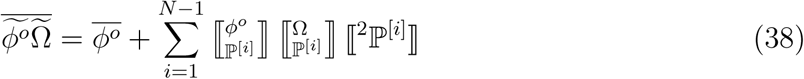

The variance term, ⟦^2^ℙ^[*i*]^⟧, can convert either of the regressions into a covariance – yielding the alternate forms of Equations 10 and 9. Combining ⟦^2^ℙ^[*i*]^⟧ with 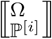, and taking the expected value, yields:

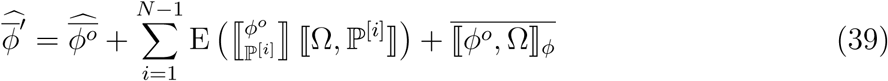

The expected value of a product in the summation can be expanded into a product of expected values and a probability covariance.

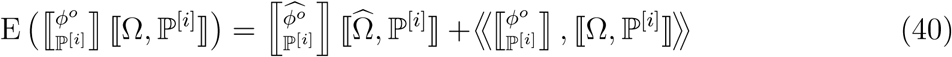

Substituting Equation 40 into Equation 39, and subtracting 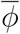 from both sides, yields Equation 10.

### Multiple traits

Consider first trait *φ*_1_. Projecting offspring phenotype into the simple basis yields:

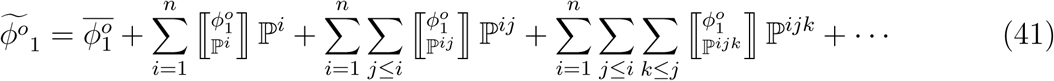

Projecting relative fitness into the conditional basis is similar, except that we also scale each term. The reason for this scaling is that we want the inner product of 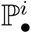 with ℙ^*i*^ to be the same as the inner product of ℙ^*i*^ with itself. We therefore divide the 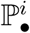 terms by the regression of the conditional term on the corresponding simple term, this is the same as multiplying terms containing 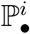 by 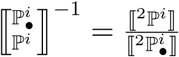, to get:

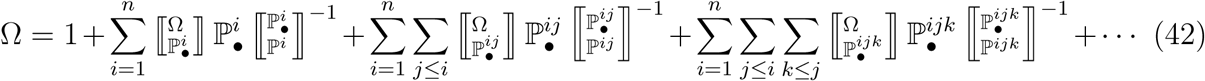

We find 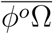 by multiplying Equations 41 and 42 and taking the mean. Keeping Equation 37 in mind, we get:

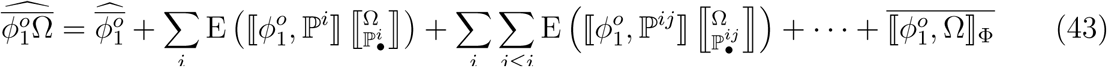

Here, 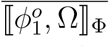 is the average covariance within sets of individuals who share the same value of *all* phenotypic traits.

Because *ϕ*^*o*^ and Ω are both random variables, so are ⟦*ϕ*^*o*^, ℙ⟧ and 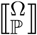. This is just saying that since each individual has a distribution of possible offspring phenotype values and a distribution of possible realized fitness values, there is a distribution of possible realized additive genetic (co)variances and a distribution of possible selection differentials. The expected values in Equation 43 can be broken up as:

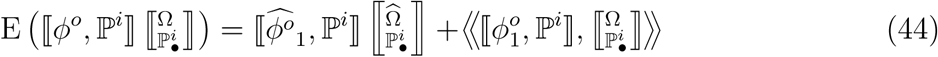

This allows us to write the expanded equation for change in each of our traits. In order to combine the equations for each trait into a single vector equation, we use the inner product and the concept of covariance between tensors.

#### The inner product

We denote the inner product of two tensors with a dot (•). For the *G*^[*i*+1]^ • *β*^[*i*]^ terms used in this paper, the inner product is always a vector (a tensor of degree 1), the elements of which are the sums of the products of the last *i* elements of *G* with all *i* elements of *β*. For example:

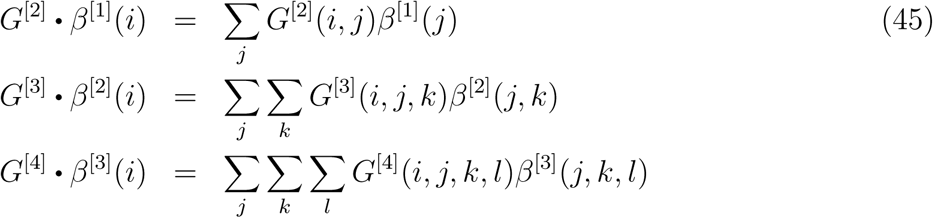

Note that in the case of G^[2]^ • *β*^[1]^, this is just standard matrix multiplication of a vector.

#### Covariance between tensors

The terms ⟪*G*^[*i*+1]^, *β*^[*i*]^⟫ in Equation 13 represent the covariance between a tensor of degree *i* + 1 (*G*^[*i*+1]^) and a tensor of degree *i* (*β*^[*i*]^). Covariance here follows the standard rule, 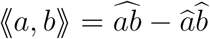, except that multiplication is replaced with the inner product, as illustrated in Equations 46.

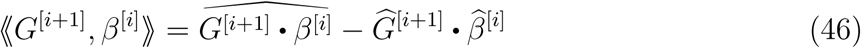

Since the degrees of the two tensors differ by 1, the result is a vector, the *i*^*th*^ element of which is the sum of all terms involving the probability covariance of 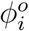 ith parental phenotype. As an illustration, ⟪*G*^[2]^, *β*^[1]^⟫ is given by:

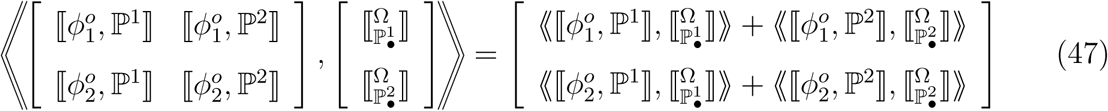

Using this notation, we can write the vector of changes in all of our traits as:

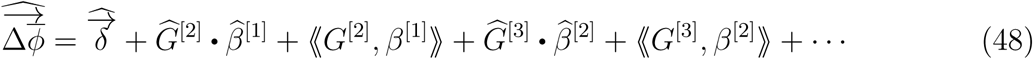

which is Equation 13.

### Numerical solutions for figures

The equilibrium distributions shown in Figures 4 through 6 were found by setting 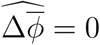 in Equation 11, with phenotypes normally distributed with variance 0.001. Since the phenotype distribution is continuous, the covariance terms were calculated using numerical integration (using scipy.integrate.quad in Python). The vector fields in Figures 10 through 13 were calculated similarly, except that each trait had a variance of 0.01.

## Supplimentary Document: Orthogonal polynomials for a population

### Orthogonality of functions

We say that two vectors are “orthogonal” if the projection of either one onto the other is zero, meaning that the inner product (or “dot product”) between them is zero. For vectors in Euclidean space, saying that the inner product is zero is equivalent to saying that the vectors are at right angles, but the concept of orthogonality extends to cases in which “right angles” has no clear meaning (Jackson, 1941).

In particular, functions can be orthogonal to one another, but here the inner product is slightly more complicated than it is for vectors. Specifically, the inner product of two functions is defined relative to a separate “weight function” (the biological meaning of which will be made clear below). The inner product of two functions, *f* (*ϕ*) and *g*(*ϕ*), with respect to a weight function, *ω*(*ϕ*), is written ⟨*f, g*⟩_*ω*_ (note the single angle brackets, distinguishing this from probability covariance). This is calculated as:

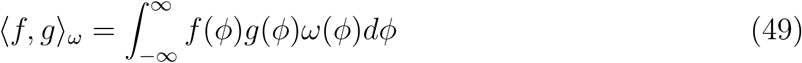

The functions *f* and *g* are said to be “orthogonal” with weight function *ω* if ⟨*f, g*⟩_*ω*_ = 0.

Different sets of orthogonal polynomials are defined by using different weight functions. For example, the Legendre polynomials, used to solve certain differential equations in physics as well as to study the evolution of continuous growth processes, use a uniform distribution on the interval (−1, 1) as their weight function.

Figure 14 illustrates one way to visualize orthogonality of functions. If either function has a mean of zero (when weighted by *ω*), then saying that the functions are orthogonal is equivalent to saying that they are uncorrelated (this will be justified below). The figure also shows the importance of the weight function, one role of which is to specify the range over which we compare the other functions.

**Figure 14:**
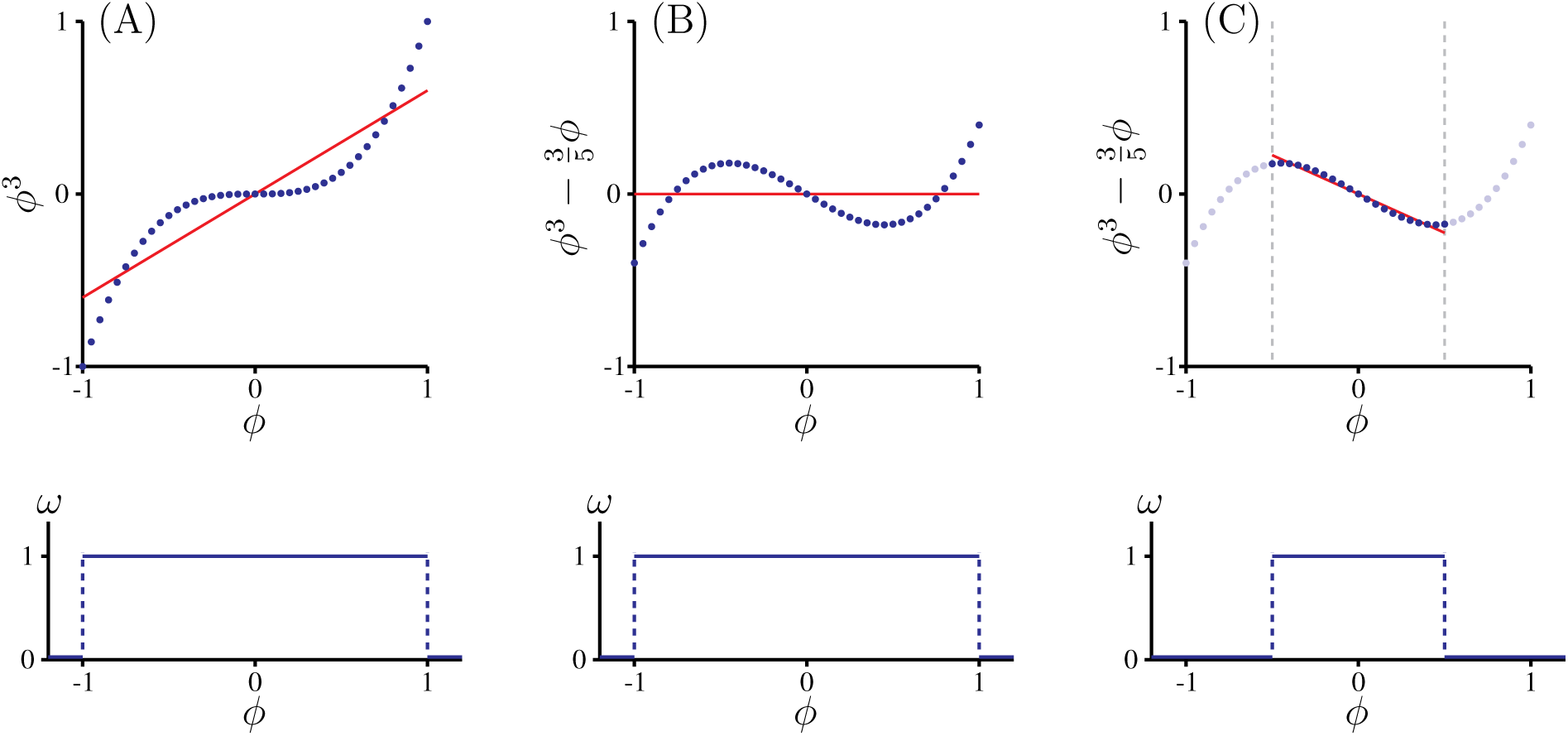
Orthogonality of functions: (A) shows that the function f = ϕ^3^ is not or- thogonal to the function g = ϕ on the interval [-1, 1] (i.e. with the weight function, ω, being 1 on the interval [-1, 1] and 0 everywhere else). The red line is the regression of ϕ^3^ on ϕ. Since 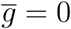 on this interval, the slope of this regression (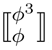) would be zero if the functions were orthogonal. (B) shows that the function 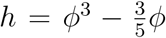 is orthogonal to g = ϕ on this interval, since the slope of the regression is zero. (C) demonstrates the importance of the weight function. If we consider the interval 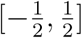 (ω = 1 on 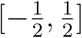 and 0 elsewhere), we find the regression slope to be clearly nonzero, meaning that h and g are not orthogonal with respect to this weight function.

The weight function need not he continuous. For example, the two blue vectors (ℙ^1^ and 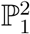) in Figure 17A are orthogonal with respect to the distribution of points (black dots) shown. Though ℙ^1^ and 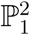 are not at right angles, the fact that they are orthogonal in the (non-Euclidean) space defined by the distribution is manifest in the fact that when we plot the points in axes defined by ℙ^1^ and 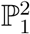, they are uncorrelated (as shown in Figure 17B). In the space of *ϕ*_1_ and *ϕ*_2_, the points are correlated; in the space of ℙ^1^ and 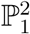, they are not. This is because ℙ^1^ and 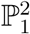 were constructed (using the methods shown below) to be orthogonal with respect to these points.

### Defining the weight function as the distribution of phenotypic variation in a population

The key to our approach is to define the weight function as the distribution of phenotypic variation in a population.

For a population of size *N*, in which individual *i* has phenotype *ϕ*_*i*_, we define the weight function over the space of all possible phenotypic values as:

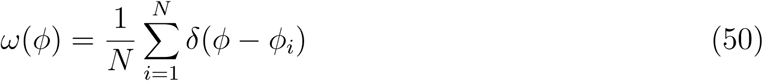

Where *δ*(*ϕ* −*ϕ*_*i*_) is the Dirac delta function, which has a value of zero everywhere except at the point *ϕ* = *ϕ*_*i*_, where its value is “infinite”. (Specifically, the magnitude of *δ*(*ϕ* − *ϕ*_*i*_) at *ϕ* = *ϕ*_*i*_ is defined such that 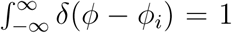. The weight function is thus basically a set of Dirac delta functions, each corresponding to one individual. We divide by *N* so that the weight function is itself a distribution that sums to 1. Figure 15 illustrates the weight function for a hypothetical population.

With this weight function, the inner product of any two functions of phenotype can be written as a sum:

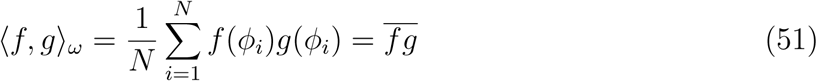

**Figure 15:**
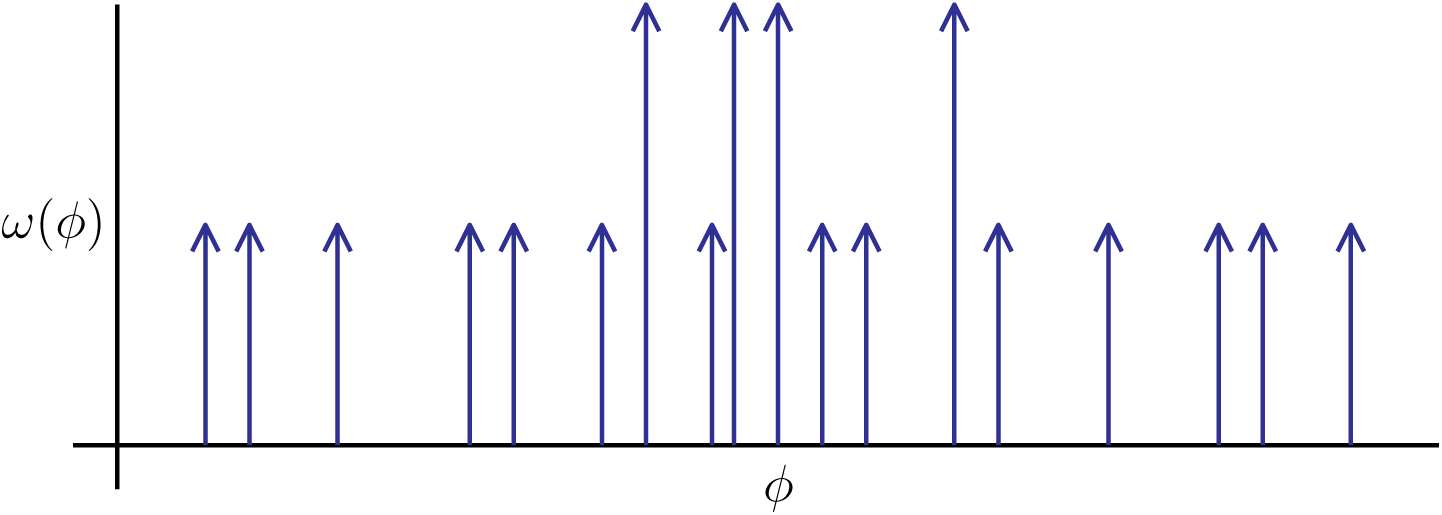
The population weight function: A population represented as a set of Dirac delta functions, each corresponding to a phenotypic value present in the population. Arrows drawn twice as long indicate two individuals with the same phenotypic value. With this weight function, the product of the functions f and g in Equation 49 is evaluated only at phenotypic values that are present in the population – turning the integral in Equation 49 into the sum in Equation 51.

If either of our functions, *f* or *g*, has a mean value of zero in the population, then the inner product becomes simply the frequency covariance:

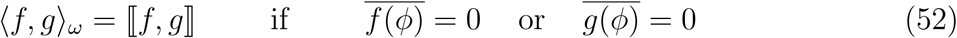

Recalling that two functions are orthogonal if their inner product is zero, this justifies our visualization of orthogonality as lack of covariation, illustrated in Figure 14.

(Note: Throughout this discussion, we denote the covariance between *f* and *g* as ⟦ *f, g*⟧, the variance of *g* as ⟦ ^2^*g*⟧, and the regression of *f* on *g* as 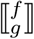.)

Note that, according to Equation 51, the inner product over a population of any function, *f*, with the constant 1 gives us 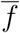, the mean value of the function in the population. This is why we set the zero^*th*^ order polynomial equal to 1:

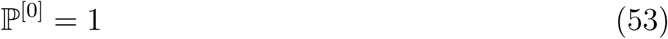

The inner product of 1 with itself is also 1, so the projection of *f* onto ℙ^[0]^ (Equation 54) is also 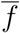.

### Constructing orthogonal functions using the Gram-Schmidt process

A key concept in constructing orthogonal polynomials is that of the projection of one vector or function onto another. A standard result is that the projection of *f* onto *g* is given by the inner product of *f* and *g* divided by the inner product of *g* with itself. Given the weight function in Equation 50, and that we set 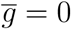, the projection of *f* onto *g* is:

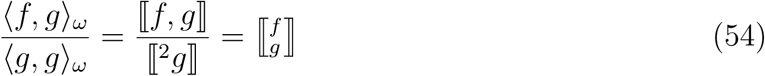

Thus, if 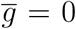, the projection of *f* onto *g* is just the slope of the (least squares linear) regression of *f* on *g*. This is why all polynomials of order ⩾ 1 are set so that their mean is zero - doing so converts inner products into moments, and projections into regressions.

Starting with first order terms, we use Gram-Schmidt orthogonalization to construct new polynomials that are orthogonal to these. Under this approach, a function for *f* independent of *g* is constructed by starting with *f* and subtracting out the projection of *f* on *g*. Using *f*_⊥*g*_ to denote a function in *f* that is orthogonal to *g*, and setting 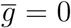 so that we can use Equation 54 for the projection, we get:

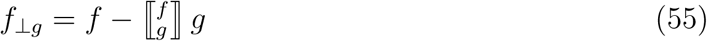

We can apply this method sequentially to construct bases in other variables that are orthogonal to these. Given a third variable, *h*, we find the function of *h* orthogonal to both *f* and *g* as:

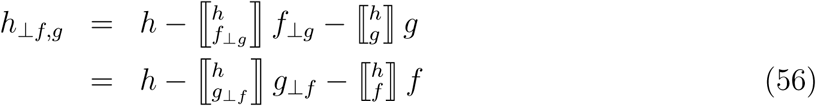

Note that there are (at least) two orthogonal bases spanning the *f* - *g* space, (*f, g*_⊥*f*_) and (*f*_⊥*g*_, *g*). Either will suffice for calculating *h*_⊥*f,g*_. What we may not do is subtract the projection of *h* on *f* and the projection of *h* on *g* (unless *f* and *g* happen to be orthogonal already).

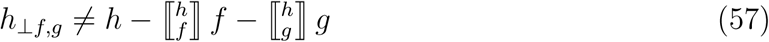

In other words, the Gram-Schmidt method works sequentially. At each step, we construct an orthogonal basis for a set of functions, then include the next function by subtracting its projection into the orthogonal basis that we already have.

### Univariate orthogonal polynomials

In this case, there is a single first order polynomial:

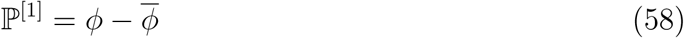

We subtract out 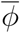 so that ℙ^[1]^ will have a mean of zero. The second order polynomial, ℙ^[2]^, is constructed so that it satisfies the following criteria:

1. The leading (first) term is (ℙ^[1]^)^2^.
2. It is orthogonal to ℙ^[1]^.
3. It has a mean of zero.

Using Equation 55, and the fact that 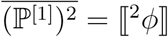, these conditions give us:

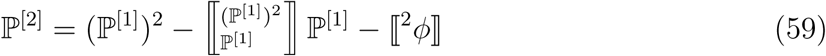

The regression of one polynomial in *ϕ* on another can be written in terms of central moments of *ϕ*. For example, to get the equation for ℙ^[2]^ used in the text, note that:

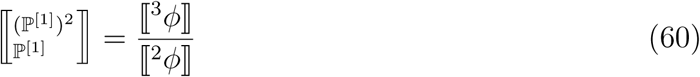

We construct the third order polynomial using the same approach. Following Equation 56, and subtracting (ℙ^[1]^)^3^ = ⟦^3^*ϕ* ⟧ so that the mean will be zero, we get:

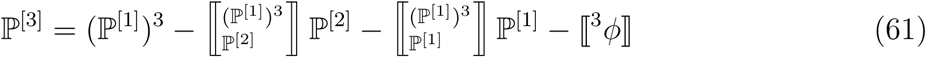

Repeating this procedure produces sequentially higher order orthogonal polynomials. The entire set of such polynomials forms the basis of a function space. Any variable can be projected into this space. Equivalently, we can write any variable of interest as a sum of terms, each containing the projection (regression) of the variable on one of the polynomials. The sum of the first *N* − 1 terms, where *N* is the number of distinct values on the *ϕ* axis, yields a function that passes through each point (or the mean, if multiple traits share the same *ϕ* value) (Figure 16).

### Multivariate orthogonal polynomials

#### First order

For the *i*^*th*^ trait, *ϕ*_*i*_, we write the first order polynomial as:

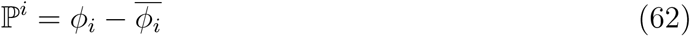

Note that there are no brackets around the superscript, indicating that this is the polynomial for trait *i*, rather than the *i*^*th*^ order polynomial.

**Figure 16:**
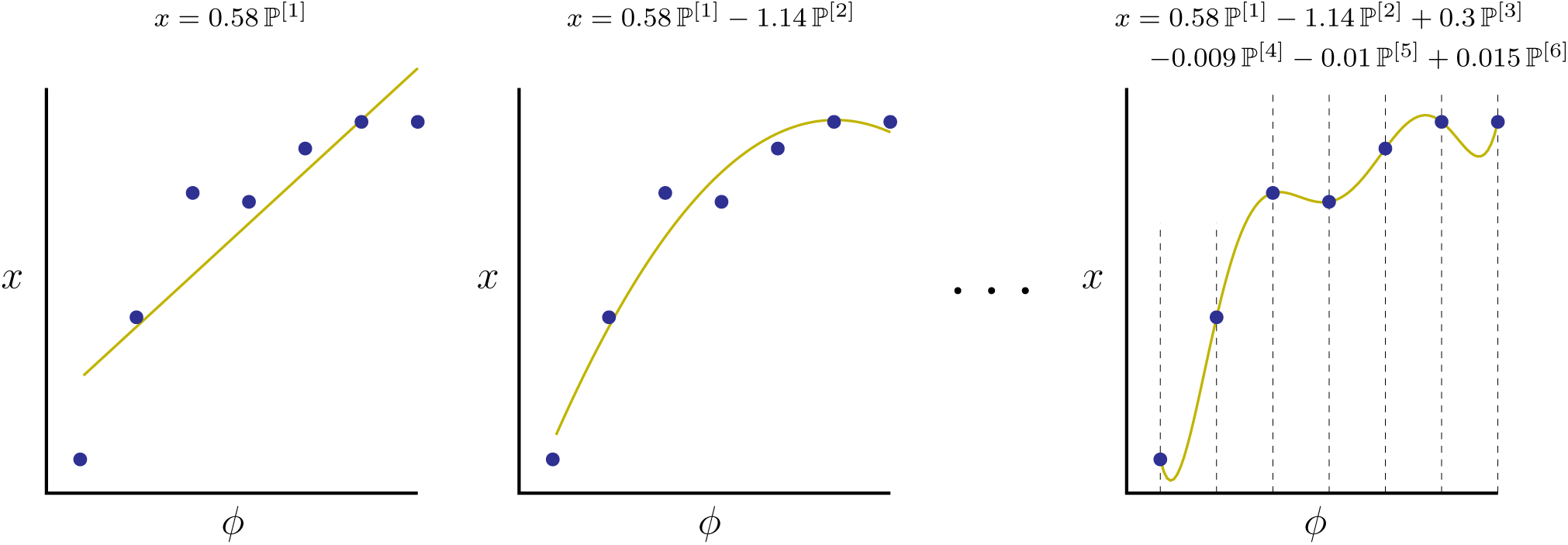
Capturing points with a series of orthogonal polynomials: Using only the first term in the polynomial expansion yields the least squares linear approximation to the data. The sum of the first and second yields the best fit quadratic function. The sum of the first N −1 terms, where N is the number of distinct values on the ϕ axis, yields a function that passes through each point (or the mean, if multiple trait values correspond to a single ϕ value). The combination of this function and the initial values of ϕ (vertical dashed lines) exactly specifies the mean of x for each value of ϕ. Note also that the first and second order coefficients do not change as we add higher order terms, as they would if we were doing an ordinary multiple regression on powers of ϕ.

For two traits, ℙ^1^ and ℙ^2^ span a two dimensional space that includes all values of *ϕ*_1_ and *ϕ*_2_, but they are not likely to be orthogonal. We thus need to construct first order orthogonal polynomials that span this space.

We will use 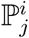 to denote a polynomial in *ϕ*_*i*_ that is orthogonal to *ϕ*_*j*_. Following Equation 55, we calculate:

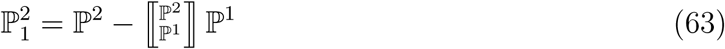

For more than two traits, we carry on following Equation 56:

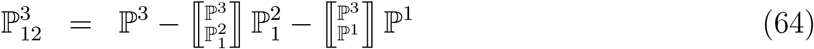

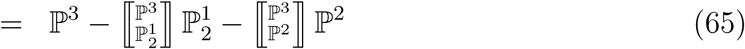

In this way, we construct a set of first order orthogonal polynomials for our set of traits. We use 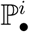 to denote the polynomial in *i* that is orthogonal to all others of the same or lower order. Thus, for the case of four traits, the polynomial in *ϕ*_2_ that is orthogonal to *ϕ*_1_, *ϕ*_3_, and *ϕ*_4_ is 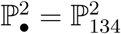.

Given another variable, *a*, the regression of *a* on ℙ^2^ is the same as the simple regression of *a* on *ϕ*_2_, while the regression of *a* on 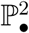 is equivalent to the *partial* regression of *a* on *ϕ*_2_ (Saville and Wood, 1991).

#### Second order

For the multivariate case, we denote second order polynomials using two superscripts, so ℙ^11^ is the second order polynomial with (*ϕ*_1_)^2^ as the leading term, ℙ^12^ is the second order polynomial with *ϕ*_1_*ϕ*_2_ as its leading term, and so forth. (If we are considering ⩾ 10 traits we can separate them with commas. For this discussion, though, we will omit the commas since we will be working with, at most, three traits. We thus write ℙ^11^ instead of ℙ^1,1^).

We want the second order polynomials to be orthogonal to all of the first order polynomials, so we apply the method from Equation 56:

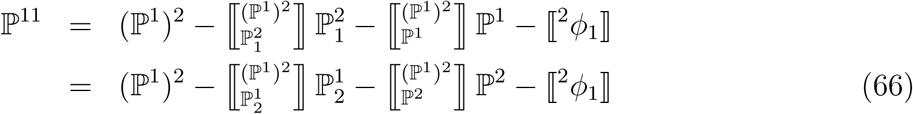

As in the univariate case, we subtract out 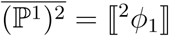 so that ℙ^11^ will have a mean of zero.

The two versions of Equation 66 correspond to two different ways of projecting (ℙ^1^)^2^ into the space spanned by the first order polynomials. Since any of the many ways of doing this will suffice, we use *𝒫*^*i*^ {*g*} to denote the projection of *g* into the space spanned by all of the *i*^*th*^ order polynomials (note that *𝒫*^[*n*]^ is orthogonal to *𝒫*^[*m*]^ for all *m* < *n*). With this notation, we can write the second order polynomials for two traits as:

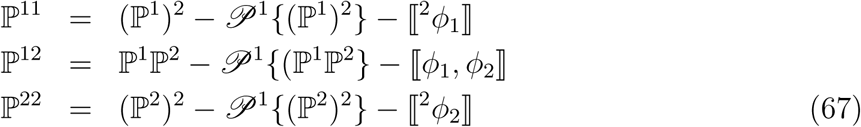

Though the second order polynomials in Equations 67 are orthogonal to all first order polynomials, they are not, in general, orthogonal to one another. To construct second order polynomials that are orthogonal to one another, we work sequentially using the Gram-Schmidt method. The polynomial in *ϕ*_1_*ϕ*_2_ that is orthogonal to ℙ^11^ and to all first order polynomials is found as:

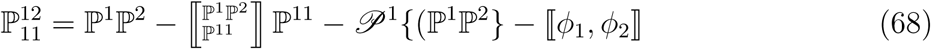

The second term on the righthand side of Equation 68 makes it orthogonal to ℙ^11^, the third term makes it orthogonal to all first order terms, and the fourth term makes it so that the mean of 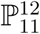 is zero.

Note that the first, third, and fourth terms on the righthand side of Equation 68 are the same at the righthand side of the equation for ℙ^12^ in Equation 67. This means that we can write Equation 68 more compactly as:

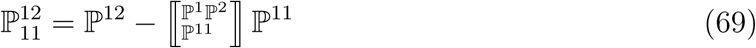

Given this term, we find the polynomial in (ℙ^2^)^2^ that is orthogonal to all other second order terms as well as all first order terms as:

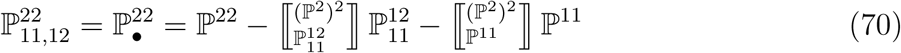

Following this approach, the remaining conditional second order terms can be written as:

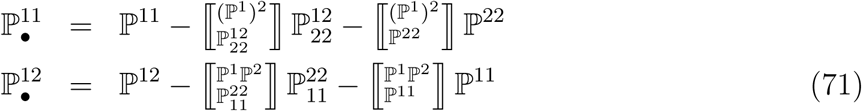

#### Biorthogonal Bases

All of the moments and regressions that we use are calculated using inner products. In fact, the principle reason that we seek orthogonal bases is that it is easy to calculate inner products in them. In a non-orthogonal basis, the standard methods for calculating inner products (and thus for calculating covariances, regressions, etc.) fail because different coordinate vectors (or functions) are correlated with one another.

Multivariate orthogonal polynomials pose a substantial challenge that does not appear in the univariate case. As shown above, finding a set of orthogonal polynomials for a multivariate distribution is relatively easy, the problem is that there are many such sets, and which one we get depends on how we choose to order our initial variables (Dunkl and Xu, 2001). Figure 17 illustrates this: The two sets of colored lines on the left represent two different bases that are orthogonal with respect to the two dimensional distribution of points shown. (Note that these bases are orthogonal with respect to a discrete distribution of points, not with respect to a uniform distribution over the space. This is why the pairs of orthogonal vectors are not at right angles. Vectors that were at right angles in the drawing would not be orthogonal with respect to the distribution of points shown). The small plots on the right of Figure 17 show the points projected into each of the bases shown on the left. The fact that these bases are orthogonal with respect to these points is manifest in the fact that there is no correlation in either of the plots on the right.

**Figure 17:**
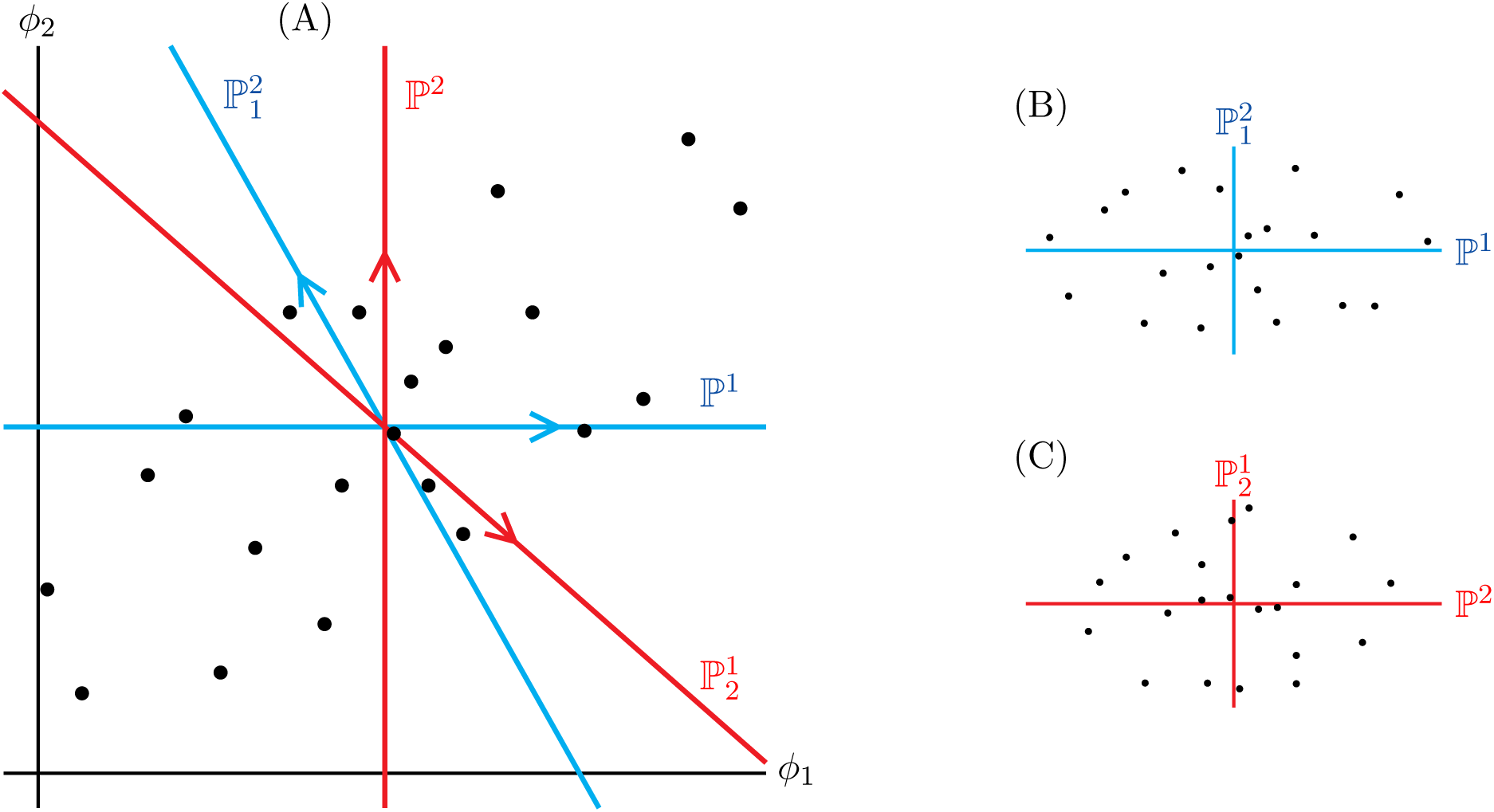
Two different orthogonal bases for a two dimensional distribution of points. Axes of the same color are orthogonal to one another with respect to this particular distribution of points. The blue axes, together, comprise one possible orthogonal basis for this space. The pair of red axes comprise another orthogonal basis. The plots on the right show the points projected into each of the bases shown on the left. The ℙ vectors are calculated as discussed in the text.

Because each of the colored bases in the figure are orthogonal with respect to the distribution, we could project fitness and offspring phenotype into any one of them and get the right answer for change in mean phenotype. Unfortunately, although the answers would be mathematically equivalent, they would be written in terms of different sets of parameters - none of which capture all of the terms that we want in our model.

For instance, if we choose to represent fitness in the blue coordinates in figure 17, we will end up with a term for the regression of fitness on *ϕ*_1_ (since ℙ^1^ points along the *ϕ*_1_ axis), but there will be no corresponding term for the regression of fitness on *ϕ*_2_, only on 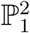 (in other words; we would have the simple regression of fitness on *ϕ*_1_, and the *partial* regression of fitness on *ϕ*_2_). The reverse is the case for the red coordinates.

None of the obvious sets of orthogonal bases in a multivariate space is satisfactory for our purposes. However, we can construct a *pair* of bases that do the job. The trick is to find a pair of bases with the property that, for each basis, each axis in it is orthogonal to all but one of the axes in the other basis. A pair of bases with this property is said to be “biorthogonal”. Formally: for two bases in an *n* dimensional space, *X* = {*x*_1_, *x*_2_, …, *x*_*n*_} and *Y* = {*y*_1_, *y*_2_, …, *y*_*n*_}, we say that *X* and *Y* are biorthogonal if:

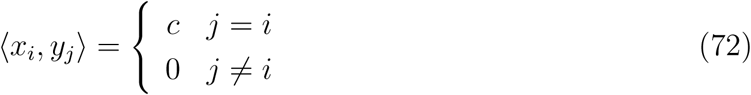

where *c* ≠ 0 is a constant. (In the special case of *c* = 1, we say that the bases are “dual”.)

Given a pair of biorthogonal bases, we can find the inner product of two functions if we first project one function into one of the pair of bases, and project the other function into the other basis (scaled by the projection of the second basis on the first). In fact, we have already done all of the work needed to get a pair of biorthogonal bases – we need only rearrange the terms that we have.

Figure 18 shows how, in two dimensions, we can construct a pair of biorthogonal bases from the two sets of orthogonal bases that we already have. We refer to the basis formed by ℙ^1^ and ℙ^2^ (which just measure the original *ϕ* values) as the simple basis, and that formed by 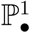 and 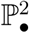 (each constructed to be orthogonal to all but one of the simple vectors) as the conditional basis.

**Figure 18:**
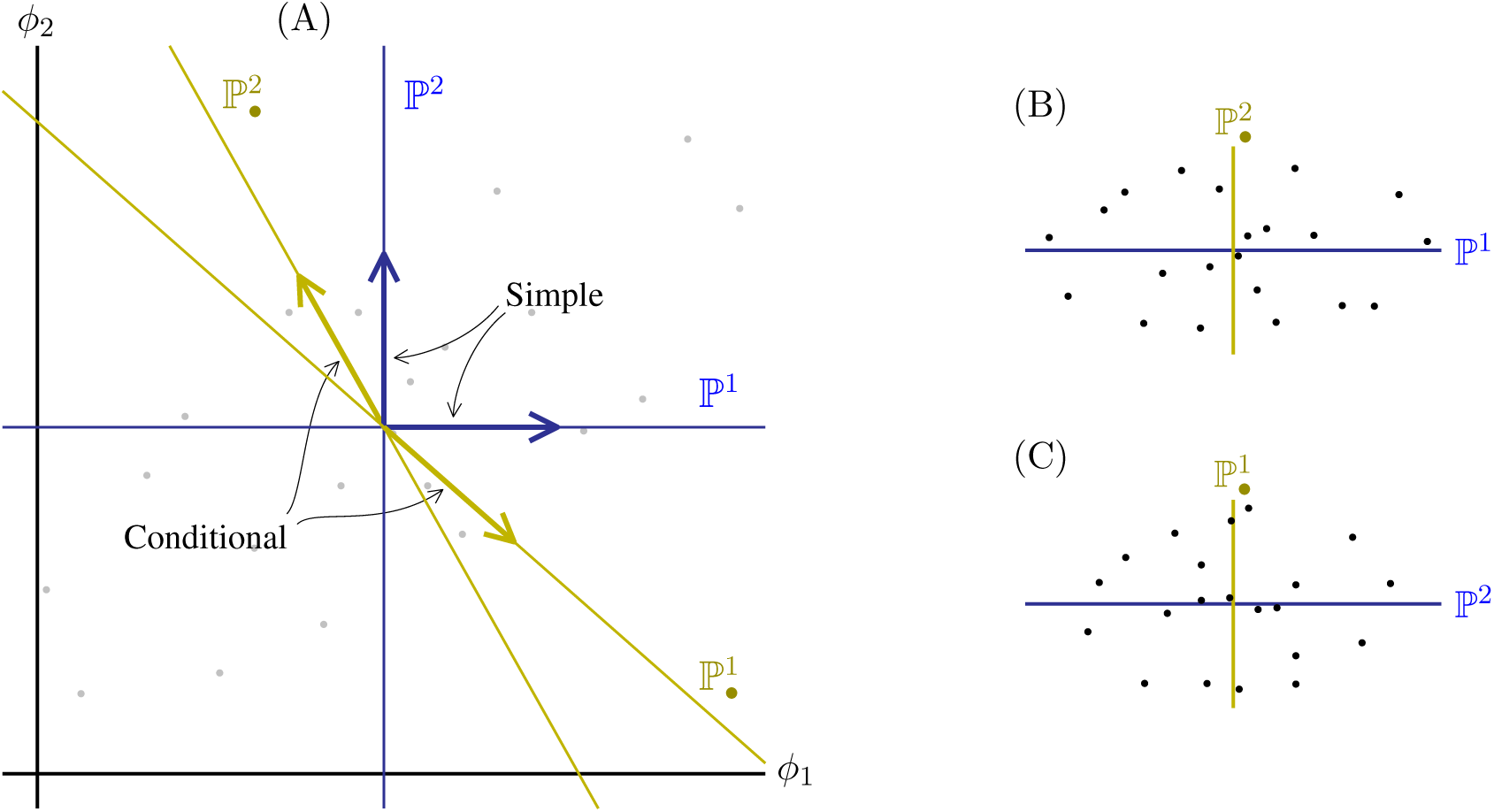
Constructing biorthogonal bases from different sets of orthogonal bases: The basis formed by vectors ℙ^1^ and ℙ^2^ (dark blue) and the basis formed by 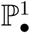 and 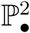 (yellow) are biorthogonal to one another. 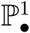 and 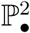 are the same as 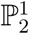 and 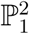 from Figure 17. In our notation, ℙ^1^ and ℙ^2^ comprise the simple basis, and 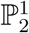 and 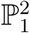 the conditional basis.

The advantage of this approach is that both bases have straightforward biological interpretations. The projection of a variable into the {ℙ^1^, ℙ^2^} basis gives the slope of the *simple* regressions of the variable on *ϕ*_1_ and *ϕ*_2_; projecting the variable into 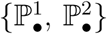 is equivalent to taking the *partial* regressions on *ϕ*_1_ and *ϕ*_2_.

Given a pair of biorthogonal bases, there is one more step necessary to calculate inner products: projections into the conditional basis (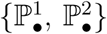 in the example) must be scaled by 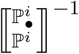 (*i.e.* we divide by the projection of the conditional basis on the simple basis (Dodson and Poston, 1991)). The reason for this is that we want the projection of 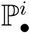 on ℙ^*i*^ to be the same as the projection of ℙ^*i*^ on itself (which is the variance in ℙ^*i*^).

To find 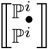, note that the variance in a function can be written as the covariance of the function with itself:

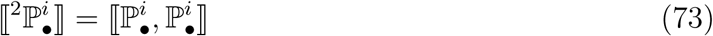

Expanding the second 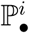 term on the right, we have:

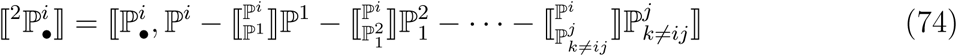

Since ⟦ *a, b* + *c*⟧ = ⟦ *a, b*⟧ + ⟦ *a, c*⟧, we can break up the righthand side of Equation 74 to yield:

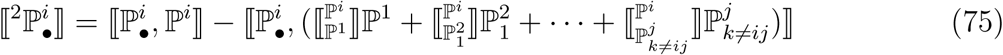

Now, 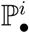 was constructed so as to be orthogonal (have zero covariance with) all of the ℙ^*j*^terms for which *j* ≠ *i*, so the second term on the righthand side of Equation 75 is zero. We thus have:

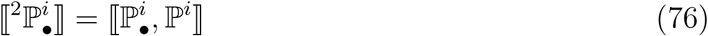

With Equation 76, we find the regression of 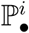 on ℙ^*i*^ to be:

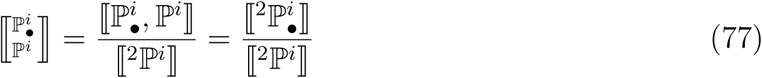

(Note that, by the same reasoning, 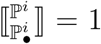. This makes sense in light of how 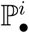 was constructed, and reminds us that the reciprocal of a regression is not just the reverse regression).

## Acknowledgements

Thanks to John Harting, for suggesting the use of orthogonal polynomials. This work was supported in part by NSF grant EF 0928772.

## References

Ardia, D. R. and E. B. Rice, 2006. Variation in heritability of immune function in the tree swallow. Evolutionary Ecology 20:491–500.

Arnold, S. J., R. Burger, P. A. Hohenlohe, B. C. Ajie, and A. G. Jones, 2008. Understanding the evolution and stability of the G-matrix. Evolution 62:2451–2461.

Beardsley, J. P., R. W. Bratton, and G. W. Salisbury, 1950. The curvilinearity of heritability of butterfat production. Journal of Dairy Science 33:93–97.

Casco, J. G., M. M. Muoz, L. S. Lpez, and C. R. Valdovinos, 2014. Genotype by environment interaction for carcass traits and intramuscular fat content in heavy iberian pigs fattened in two different free-range systems. Spanish Journal of Agricultural Research 12:388–395. URL http://revistas.inia.es/index.php/sjar/article/view/4840.

Charmantier, A. and D. Garant, 2005. Environmental quality and evolutionary potential: lessons from wild populations. Proceedings of the Royal Society B: Biological Sciences 272:1415–1425.

Dodson, C. T. J. and T. Poston, 1991. Tensor geometry: the geometric viewpoint and its uses. Second Edition. Springer-Verlag, Berlin.

Dunkl, C. F. and Y. Xu, 2001. Orthogonal polynomials of several variables. Cambridge Univ. Press, New York.

Fuerst-Waltl, B., J. Solkner, A. Essl, I. Hoeschele, and C. Fuerst, 1998. Non-linearity in the genetic relationship between milk yield and type traits in Holstein cattle. Livestock Production Science 57:41–47.

Gifford, D. R. and J. S. F. Barker, 1991. The nonlinearity of offspring-parent regression for total sternopleural bristle number of drosophila melanogaster. TAG Theoretical and Applied Genetics 82:217–220. URL http://dx.doi.org/10.1007/BF00226216.10.1007/BF00226216.

Gimelfarb, A., 1986. Offspring-Parent Genotypic Regression: How Linear Is It? Biometrics 42:67–71.

Gimelfarb, A. and J. H. Willis, 1994. Linearity Versus Nonlinearity of Offspring-Parent Regression: an Experimental Study of Drosophila melanogaster. Genetics 138:343–352.

Griswold, C. K., R. Gomulkiewicz, and N. Heckman, 2008. Hypothesis testing in comparative and experimental studies of function-valued traits. Evolution 62:1229–1242.

Hansen, T., C. Pelebon, and D. Houle, 2011. Heritability is not evolvability. Evolutionary Biology 38:258–277.

Heywood, J. S., 2005. An exact form of the breeder’s equation for the evolution of a quantitative trait under natural selection. Evolution 59:2287–2298.

Hoffmann, A. A. and J. Merila, 1999. Heritable variation and evolution under favourable and unfavourable conditions. Trends in Ecology and Evolution 14:96 –101.

Holloway, G. J., S. R. Povey, and R. M. Sibly, 1990. The effect of new environment on adapted genetic architecture. Heredity 64:323–330.

Houle, D., 1992. Comparing evolvability and variability of quantitative traits. Genetics 130:195–204.

Husby, A., M. E. Visser, and L. E. B. Kruuk, 2011. Speeding up microevolution: The effects of increasing temperature on selection and genetic variance in a wild bird population. PLoS Biol 9:e1000585.

Jackson, D., 1941. Fourier series and orthogonal polynomials. Mathematical Association of America, Oberlin, OH.

Kirkpatrick, M., D. Lofsvold, and M. Bulmer, 1990. Analysis of the Inheritance, Selection and Evolution of Growth Trajectories. Genetics 124:979–993.

Kole, P. and A. Saha, 2013. Studies on variability and heritability for different quantitative characters in fenugreek under different environments. J. Plant Breed. Crop Sci. 5:224–228.

Konarzewski, M., A. Ksiazek, and I. B. Lapo, 2005. Artificial selection on metabolic rates and related traits in rodents. Integrative and Comparative Biology 45:416–425. URL http://icb.oxfordjournals.org/content/45/3/416.abstract.

Lande, R. and S. J. Arnold, 1983. The measurement of selection on correlated characters. Evolution 37:1210–1226.

Lynch, M. and B. Walsh, 1998. Genetics and Analysis of Quantitative Traits. Sinauer Associates, Sunderland, MA.

McElroy, T. C. and W. J. Diehl, 2001. Heterosis in two closely related species of earthworm (*Eisenia fetida* and *E. andrei*). Heredity 87:598–608.

Melo, D. and G. Marroig, 2015. Directional selection can drive the evolution of modularity in complex traits. Proc. Nat. Acad. Sci. USA 112:470–475.

Merila, J. and B. C. Sheldon, 2001. Avian quantitative genetics. chap. 4, P. 179, in V. N. Jr., ed. Current Ornithology, Vol 16. Kluwer/Plenum, New York.

Meyer, H. H. and F. D. Enfield, 1975. Experimental evidence on limitations of the heritability parameter. TAG Theoretical and Applied Genetics 45:268–273. URL http://dx.doi.org/10.1007/BF00831900.10.1007/BF00831900.

Meyer, K. and M. Kirkpatrick, 2005. Up hill, down dale: quantitative genetics of curvaceous traits. Philos Trans R Soc Lond B 360:1443–1455.

Nishida, A., 1972. Some characteristics of parent-offspring regression in body weight of Mus musculus at different ages. Canadian Journal of Genetics and Cytology 14:292–303.

Pletcher, S. D. and C. J. Geyer, 1999. The Genetic Analysis of Age-Dependent Traits: Modeling the Character Process. Genetics 153:825–835.

Rice, S. H., 2004. Evolutionary theory: mathematical and conceptual foundations. Sinauer Associates, Sunderland, MA.

Rice, S. H., 2008. A stochastic version of the Price equation reveals the interplay of deterministic and stochastic processes in evolution. BMC Evolutionary Biology 8:262.

Rice, S. H., 2012. The place of development in mathematical evolutionary theory. Journal of Experimental Zoology Part B: Molecular and Developmental Evolution 318:480–488. URL http://dx.doi.org/10.1002/jez.b.21435.

Rice, S. H. and A. Papadopoulos, 2009. Evolution with Stochastic Fitness and Stochastic Migration. PLoS ONE 4:e7130.

Rice, S. H., A. Papadopoulos, and J. Harting, 2011. Stochastic processes driving directional evolution. in P. Pontarotti, ed. Evolutionary Biology. Springer-Verlag, Berlin (In press).

Ryan, P. G., 2001. Morphological heritability in a hybrid bunting complex: *Neospiza* at inaccessible island. The Condor 103:429–438.

Saastamoinen, M., J. E. Brommer, P. M. Brakefield, and B. J. Zwaan, 2013. Quantitative genetic analysis of responses to larval food limitation in a polyphenic butterfly indicates environment- and trait-specific effects. Ecology and Evolution 3:3576–3589. URL http://dx.doi.org/10.1002/ece3.718.

Saville, D. J. and G. R. Wood, 1991. Statistical methods: the geometric approach. Springer-Verlag, New York.

Shimizu, H. and T. Awata, 1979. On the linearity of heritability and genetic correlation for juvinile body weight and weight gain in meat-type chickens. J. Fac. Agr. Hokkaido Univ. 59:333–345.

Steppan, S. J., P. C. Phillips, and D. Houle, 2002. Comparative quantitative genetics: evolution of the Gmatrix. Trends Ecol. Evol 17:320–327.

Stinchcombe, J. R., Function-valued traits working group, and M. Kirkpatrick, 2012. Genetics and evolution of function-valued traits: understanding environmentally responsive phenotypes. Trends Ecol. Evol. 27:637–647.

Wilson, A. J., J. M. Pemberton, J. G. Pilkington, D. W. Coltman, D. V. Mifsud, T. H. Clutton-Brock, and L. E. B. Kruuk, 2006. Environmental coupling of selection and heritability limits evolution. PLoS Biol 4:e216. URL http://dx.doi.org/10.1371/journal.pbio.0040216.

